# A Tandem Repeat Atlas for the Genome of Inbred Mouse Strains: A Genetic Variation Resource

**DOI:** 10.1101/2025.05.23.655792

**Authors:** Wenlong Ren, Weida Liu, Zhuoqing Fang, Egor Dolzhenko, Ben Weisburd, Zhuanfen Cheng, Gary Peltz

## Abstract

Tandem repeats (TRs) are a significant source of genetic variation in the human population; and TR alleles are responsible for over 60 human genetic diseases and for inter-individual differences in many biomedical traits. Therefore, we utilized long-read sequencing and state of the art computational programs to produce a database with 2,528,854 TRs covering 39 inbred mouse strains. As in humans, murine TRs are abundant and were primarily located in intergenic regions. However, there were important species differences: murine TRs did not have the extensive number of repeat expansions like those associated with human repeat expansion diseases and they were not associated with transposable elements. We demonstrate by analysis of two biomedical phenotypes, which were identified over 40 years ago, that this TR database can enhance our ability to characterize the genetic basis for trait differences among the inbred strains.

## INTRODUCTION

Tandem Repeats (TR) are highly polymorphic sequences that contain repeated copies of a short motif, which are distributed throughout the genome ^1,2^. Over 15 years ago, it was postulated that TR alleles could be responsible for a significant percentage of the un-identified genetic factors (i.e., ‘missing heritability’) that determine many human trait differences and disease susceptibilities ^3^. Recently developed methods for characterizing TR alleles ^4,5,1^ has enabled TR allelic effects to be characterized. Consistent with their potential to contribute to ‘*missing heritability’*, TRs cover 6 to 8% of the human genome ^6,7^; TR expansions have been associated with 65 neurological and 14 neuromuscular conditions, which include Huntington’s disease and fragile X syndrome ^8,9,10^; and most TR expansion diseases were initially characterized over the last 10 years. Genetic association studies found that TR alleles: had a strong association with multiple human phenotypes (height, hair morphology, biomarkers, etc.) ^4^; influenced 58 complex traits; modulated the expression or splicing of a nearby gene (n=18); and were the largest contributors to glaucoma and colorectal cancer risk ^5^. Consistent with their association with brain diseases, TR alleles affect the expression and splicing of many mRNAs in brain, and brain phenotypes (i.e., cortical surface area) ^11^. Characterization of human TR alleles has also uncovered new regulatory mechanisms and a potential new treatment for a human disease. TR alleles within cis-regulatory elements can affect gene expression by forming structures that alter transcription factor binding ^12^. Repeat expansions can lead to protein synthesis without AUG initiation codons that occurs from multiple reading frames and in multiple directions (i.e., repeat associated non-AUG (or RAN) translation) ^13^. This provides a mechanism for some TR expansion diseases. (e.g., myotonic dystrophy type 2 (DM2)) ^14^. Also, RAN-associated TR expansions form hairpin structures that activate double stranded RNA-dependent protein kinase (PKR), which impairs the translation of most proteins but increases RAN translation. Moreover, treatment with a widely used diabetes drug (metformin), which decreased RAN translation, improved the behavioral phenotypes in a mouse model of frontotemporal dementia ^15^.

Characterization of the genetic architecture of murine models for human diseases has provided insight into many human diseases ^16^. We recently demonstrated that characterization of structural variants in the mouse genome facilitated the identification of a causative genetic factor for a murine lymphoma model that was first described over fifty years ago ^17^. Given the importance of TR alleles to human disease, we used high-fidelity long-read genomic sequencing and new computational tools to comprehensively characterize TR alleles in 39 inbred strains. We observed that there was significant diversity among the TRs in different mouse strains, and there were significant differences in the properties of the TRs present in mice and humans. We demonstrate the importance of TR alleles for genetic discovery by analyzing two biomedical phenotypes, which were characterized in inbred strains over 40 years ago, but the causative genetic factors for them were not previously identified.

## RESULTS

### Genomic sequencing and TR genotyping

Long-read (genomic) sequencing (LRS) was performed on 40 inbred strains (30-fold genome coverage per strain) using a PacBio Revio platform equipped with the HiFi system ^18^ (**Table S1**). Perfect tandem repeats (TRs) in their genomes were identified using the pipeline shown in **Figure 1**. In brief, the sequence data was first analyzed using the Tandem Repeat Genotyping Tool ^19^, and then variation clustering was performed ^20^ for 39 strains using C57BL/6 as the reference sequence (GRCm39) to produce a catalog with 3,494,901 TRs. Since they are commonly used in genetic models, we separately report on the TRs present in the 35 classical inbred strains and those in all 39 sequenced strains, which includes the four wild derived strains (CAST, SPRET, MOLF, WSB). After removing non-polymorphic (i.e., with alleles identical to the reference genome) and potentially mosaic TRs, the final dataset consisted of 2,528,854 (or 1,819,293) TRs in the 39 (or 35 classical) inbred strains (Figure1; **Table S2**). The percentage of TR genotypes in this database for the 39 (or 35) inbred strains was 99.3% (or 99.6%), which indicates that there is an extremely low rate (0.4-0.7%) of absent genotypes.

**Figure 1.**
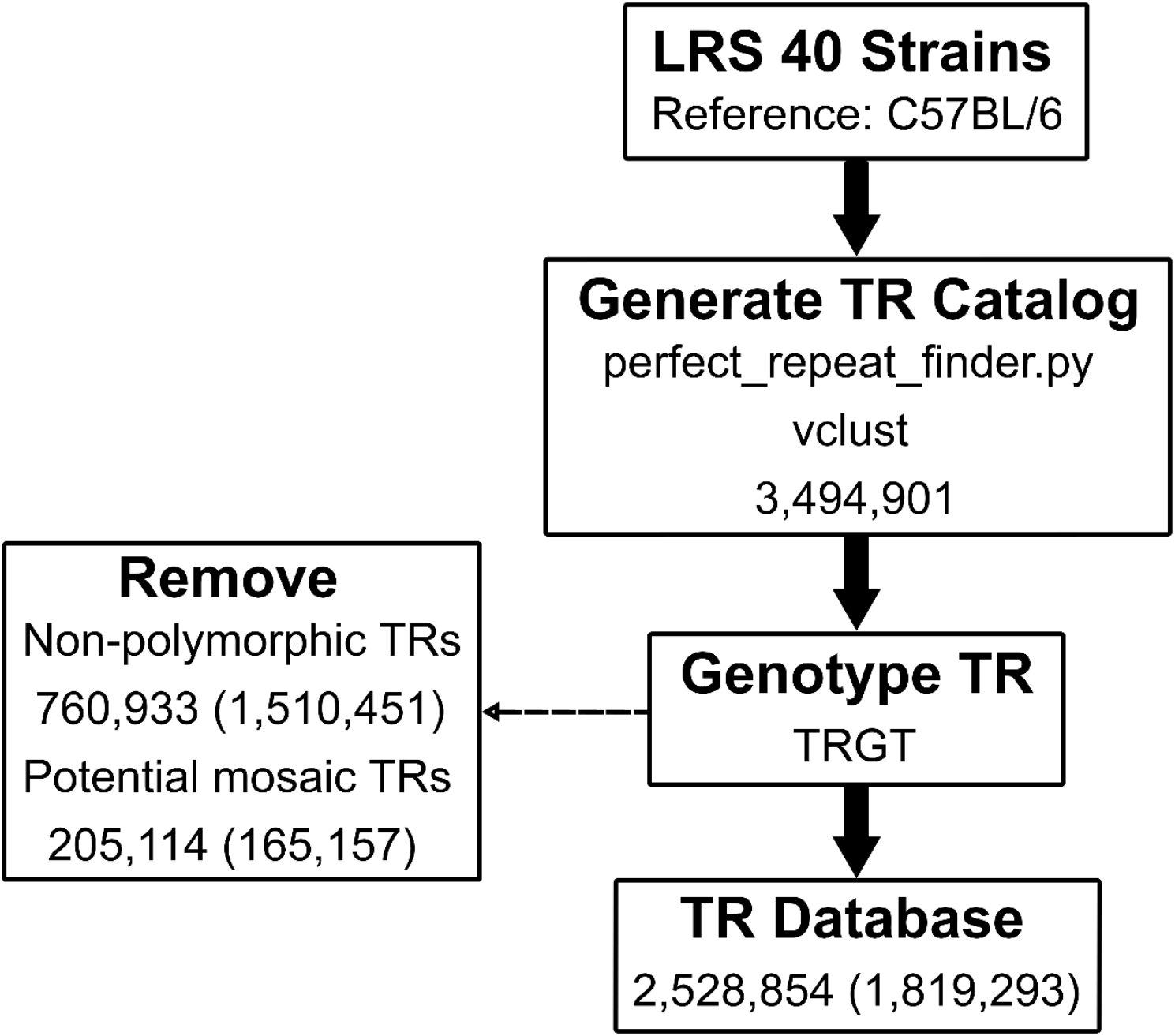
Overview of the pipeline used to analyze the genomic sequences of 40 inbred mouse strains to generate the TR database. Long Read Sequencing (LRS) was performed on 40 inbred strains, and C57BL/6 was used as the reference sequence. The programs used to generate the TR catalog and for TR genotyping are shown. The TRs in all 39 (or 35 classical) inbred strains were merged. The TRs in all strains matching the reference sequence (i.e., non-polymorphic TRs) or where heterozygous alternative alleles (i.e., potential mosaic TRs) were detected were removed. A TR database with 2,528,854 (1,819,293) was established. The numbers within parenthesis indicate the number of TRs present in the 35 classical inbred strains.

### TR Characterization

More TRs were found in the four wild-derived inbred strains than in the 35 classical inbred strains, and three wild-derived strains (SPRET, CAST and MOLF) had a particularly high number of strain-unique TRs. For example, SPRET mice had 2.5 times more TRs (n=1,773,873) than were found in any of the 35 classical inbred strains; and SPRET mice had the highest number of (n=286,391) strain-unique TRs. Most minor TR alleles are shared by 1 to 3 strains (**Figure S1**) and the number of TRs decreased as the number of strains sharing a minor TR allele increased (**Figure 2**). Among the 35 classical inbred strains: CE mice had the highest number of TRs (n=728,155); the strains most closely related to the C57BL/6 reference strain (B10J, n=84,069; and B10D2, n=89,490) had the fewest; and KK mice had the highest number of strain-unique TRs (n=21,199), which is ∼48 times greater than was found in B10D2 mice. CE (n=18,683), SMJ (n=16,207), and TallyHo (n=14,340) mice also had many strain-unique TRs (**Table S3**). There was an average of 6.52 alleles per TR among the 35 classical inbred strains.

**Figure 2.**
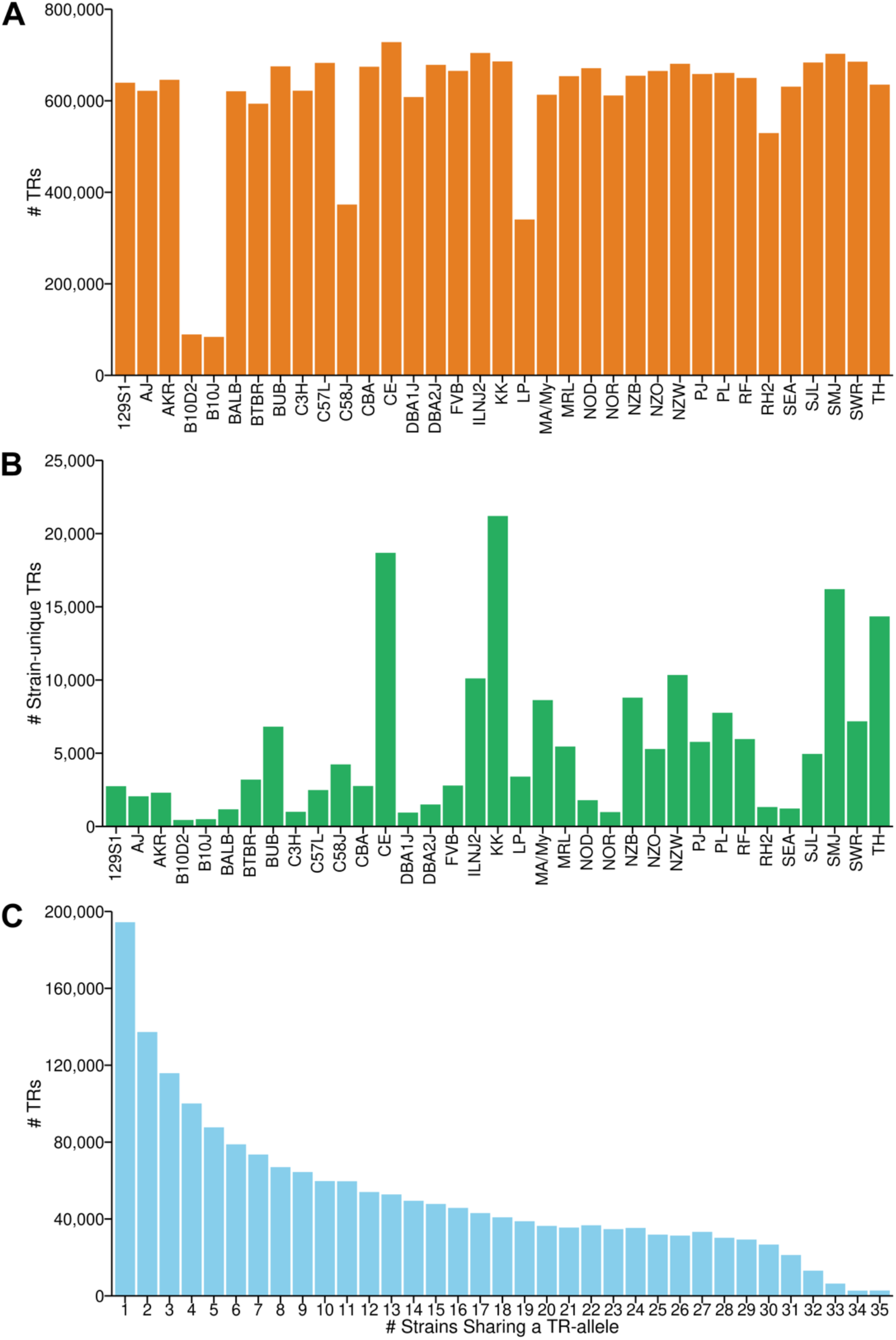
The distribution and characteristics of TRs in 35 classical inbred strains. (A and B) The total number of TRs (A) and the number of strain-unique TRs (B) are shown for each strain. Four strains (CE, KK, SMJ, and TallyHo (TH)) possess a greater number of strain-unique TRs. (C) The number of TRs where a minor allele is shared by the indicated number of strains is shown. Most of the minor TR alleles are shared by 1-3 strains.

When their genomic locations were analyzed, most murine TRs were intergenic (n=1,428,904 for the 39 strains) or intronic (n=985,616), which is consistent with the distribution of human TRs ^10^. However, some mouse TRs were in coding regions: 77,318 (or 53,990) were exonic; 31,140 (or 21,563) were within 3’ UTRs; 5,876 (or 3810) were in 5’ UTRs; and 2,539 (or 1847) TRs were near transcriptional start sites (TSS) in all 39 (or 35 classical) inbred strains (**Figures 3A and S2A**). TRs with motif lengths of 2, 3, or 4 are the most abundant type of TR. The 1,901,163 (or 1,573,675) TRs with a motif length of two present in all 39 (or 35 classical) inbred strains significantly surpasses the number of TRs with other motif lengths. In contrast, TRs with motif lengths greater than 6 are much less common. Human TR alleles can be highly polymorphic ^2^, but most murine TR alleles have a single motif: 1,947,184 (or 1,283,601) single motif alleles are present in all 39 (or 35 classical) inbred strains. However, murine TR alleles containing 2 to 3 motifs are also relatively frequent (>100,000 of each type), but the number of TR alleles with 4 or more distinct motifs was substantially reduced (**Figures 3B-3C and S2B-S2C**).

**Figure 3.**
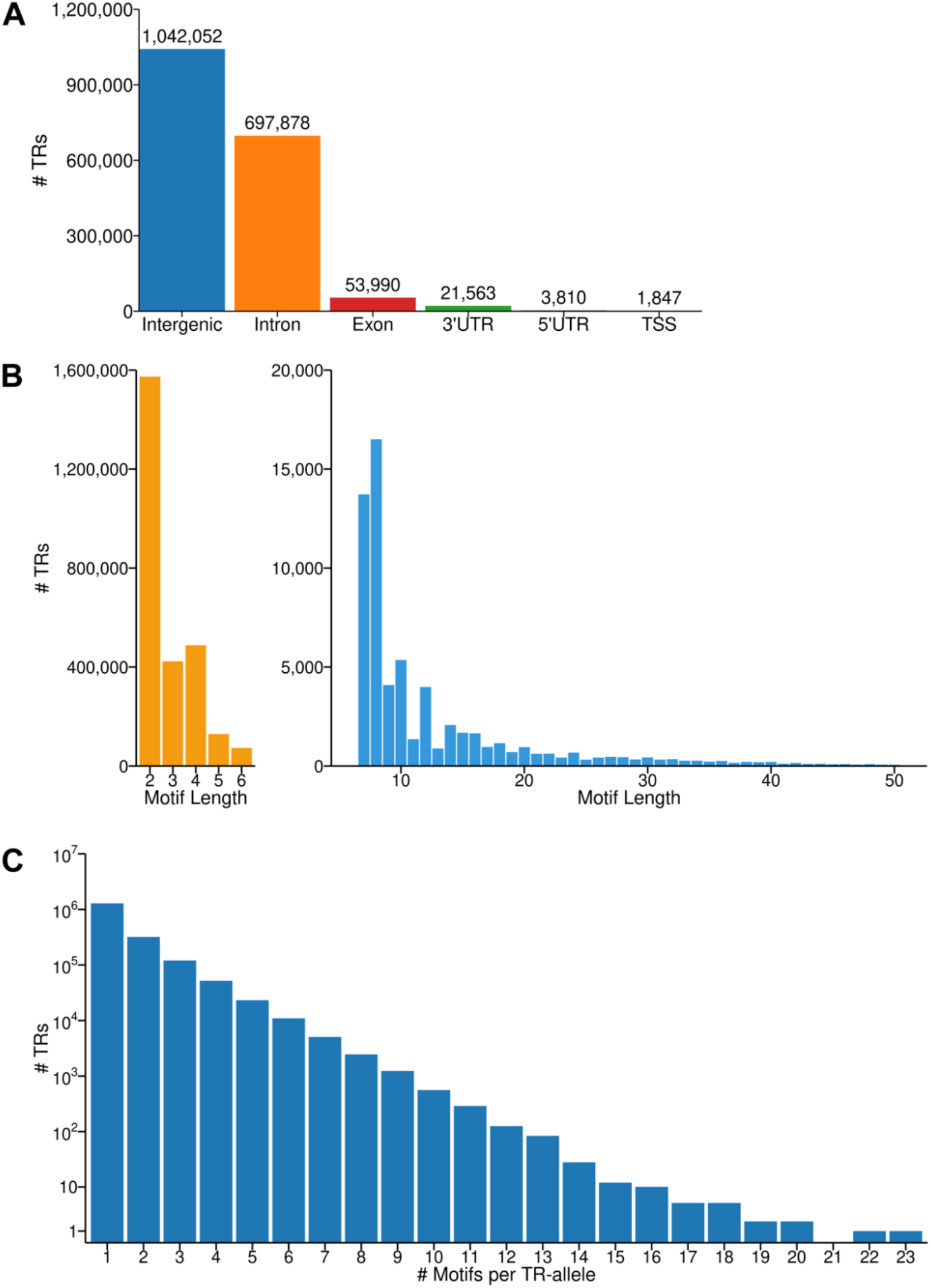
The genomic distribution and properties of TRs in the 35 classical inbred strains. (A)The distribution of TRs in different types of genomic regions. (B)The number of TRs with different motif lengths. Most TRs are <7 bp (left), while TRs with motifs >6 bp are rarer (right). (C)The number of TRs with alleles with the indicated number of motifs. The Y-axis is log_10_ transformed.

Thirty TRs, which included those with single-motif and compound-motif repeats, were validated by PCR-amplification and analysis of amplicon size or (when needed) band sequencing in 6 inbred strains (AJ, B10J, CBA, NOD, TH and C57BL/6J), with C57BL/6J serving as the reference (**Table S4**). For example, a TR at chr6:29099453-29099501 has only a single GT motif [AT(GT)_9_G] relative to the [AT(GT)_23_G] allele in the reference strain, which represents a fourteen-unit contraction in AJ. Similarly, a compound TR at chr1:81132699-81132743 has a TallyHo allele [C(TCTCTG)_3_(TC)_6_] while C57BL/6 has a [C(TCTCTG)_4_(TC)_10_] allele, which reflects losses of one TCTCTG and four TC motifs in the TallyHo genome. All thirty of these TR loci yielded the expected amplicons from each of the 5 strains for the predicted alleles. This result confirms the accuracy of our TR database, which results from the generation of high-fidelity genomic sequence, abundant sequence coverage, and from the robustness of the computational pipeline used for its construction.

### TRs in murine homologues of human repeat expansion disease causing genes

Since over 65 human diseases results from TR expansions ^8,9,10^, we characterized the TRs present in murine homologues of human TR expansion disease genes. TR alleles that affected coding regions were identified within 31 of these genes. While most were within 3’ or 5’ UTRs, only 6 genes had exonic TRs (**Table S5)**. Although human diseases appear only when many copies of a TR are present (e.g. 50-11,000 repeats for DM2) ^10^, the number of repeats in murine exonic TR alleles only differed by <2 from the reference strain. Moreover, these murine TR alleles inserted (or removed) 1 or 2 amino acids of the protein sequence and did not disrupt the reading frame of the encoded protein. These results indicate that unlike human TRs, the number of copies of a TR in the murine genome is tightly controlled, and pathologic conditions resulting from TR expansions are unlikely to develop in the inbred strains.

### Linkage disequilibrium (LD) analysis

LD decay analysis for 35 inbred strains was performed using alleles generated from four different sets of genetic variants: (i) 220K structural variants (SVs), (ii) 1.8M TRs, and (iii) 21M or (iv) 220K single nucleotide polymorphisms (SNPs). A selected subset of SNP alleles was analyzed, which was equal the number of SV alleles analyzed, to ensure that any differences did not result from evaluation of different numbers of genetic variants. While the maximum LD values (r^2^) calculated for the 21M SNP (0.81), 220K SNP (0.76) and 1.8M TR (0.85) datasets were similar; the (r^2^) calculated for the 220K SV was 0.49. The calculated distance where the LD dropped to half of its maximum value (half decay point) were: 133 kb for the 21M SNPs, 177 kb for the 220K SNPs, 291 kb for the 220K SVs, and 0.1 kb for the 1.8M TRs (**Figure 4**). There were notable differences in LD decay patterns among the different types of variants. SNPs exhibited a relatively moderate degree of LD decay, they had a higher initial level of LD, and the decay distance ranged from 133 to 177 kb. SVs had a lower initial level of LD (0.49) but had a more extended decay distance (291 kb). While TRs had the highest initial level of LD (0.850), they had the most rapid rate of decay; the half maximal LD decay occurred within only 0.1 kb.

**Figure 4.**
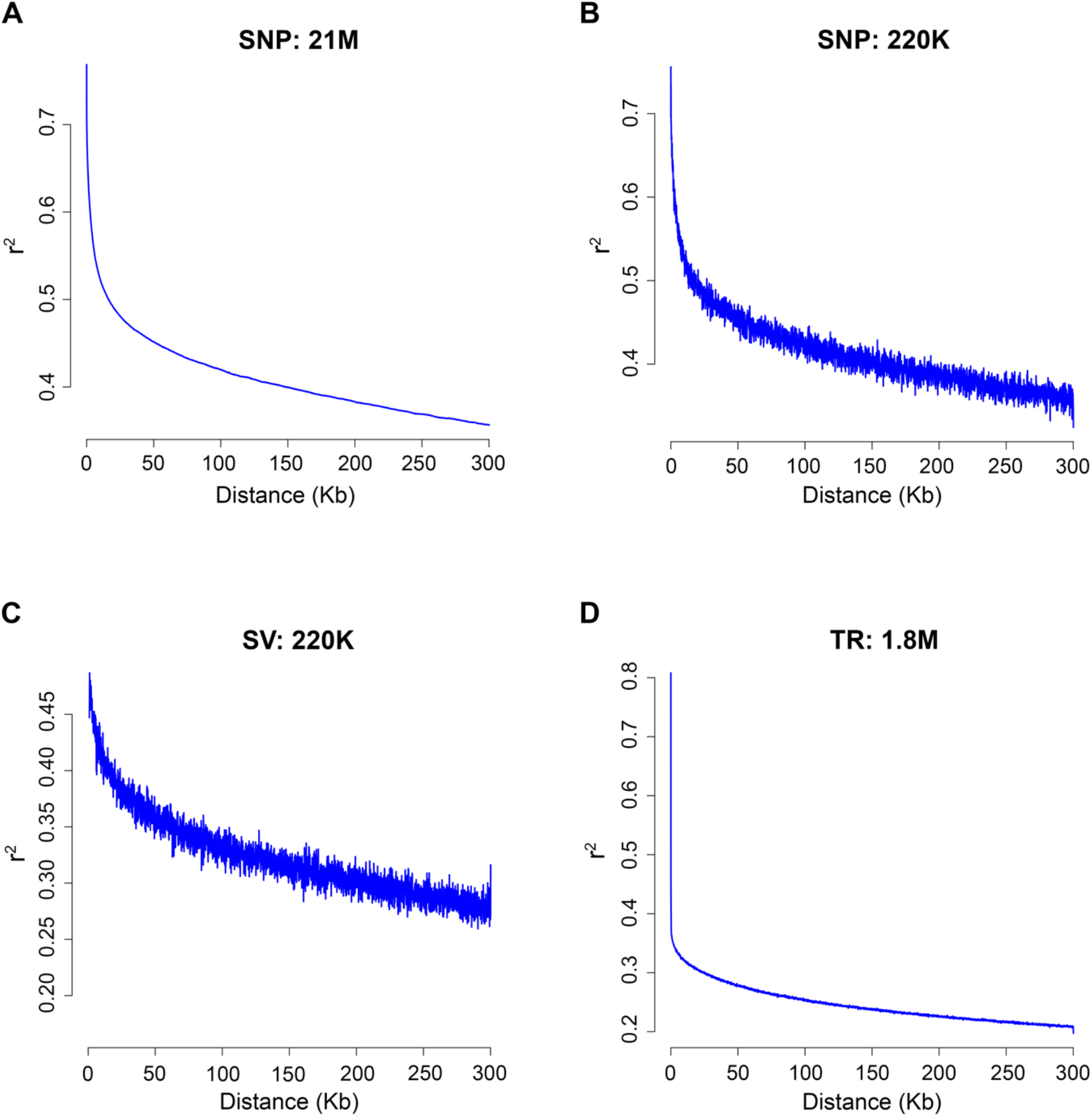
Linkage disequilibrium (LD) decay patterns across different types of genetic variants in 35 inbred strains. The LD patterns were calculated using: (A)21 million SNPs (B)220K SNPs (C)220K structural variants (SVs) (D)1.8 million tandem repeats (TRs) The y-axis represents LD values (r^2^), and the x-axis indicates physical distance (kb). The maximum LD values are 0.811, 0.756, 0.487, and 0.850, with LD decaying to half of these values at 133 kb, 177 kb, 291 kb, and 0.1 kb, respectively.

### Murine TRs are not preferentially located near transposable elements (TEs)

In the human genome, TRs occur in regions with TEs, especially the TEs containing Alu elements; but human TRs are not associated with LINE-1 insertions ^21,22^. Alu elements are not present in the mouse genome; but murine LINE-1 (18% of the genome), B1 (2.7%) and B2 (2.4%) TEs are abundant ^23^. Therefore, we investigated whether murine TRs were located near LINE-1 elements. Analysis of the TRs in the 35 classical inbred strains revealed that 68,744 TRs (3.78% of the total) were entirely within LINE-1 elements; 2,332 TRs (0.13%) overlapped with LINE-1 sequences; and 1,435 TRs (0.08%) were proximal (i.e., located within 80 bp) to LINE-1 elements. However, 1,744,506 TRs (96% of the total) were >200 bp away from a LINE-1 element. A similar distribution was observed when TRs in 39 strains were examined:, 94,694 TRs (3.74%) were within LINE-1 elements; 2,517 TRs (0.10%) overlapped with LINE-1 elements; 2,012 TRs (0.08%) were proximal to LINE-1 elements; and 2,426,539 TRs (96%) were not located near a LINE-1 element. Although ∼23% of the mouse genome consists of TEs, less than 4% of murine TRs are located in or near a TE.

### Phylogenetic analyses

The phylogenetic trees constructed for 40 inbred strains using three types of genetic variants [SNP, structural variant (SV), and TR alleles] generally reflected the known evolutionary and phylogenetic relationships among the strains (**Figure S3**). As examples, the four wild-derived strains (WSB, MOLF, SPRET, and CAST) were separated from the classical inbred strains; the NZW, NZO, and NZB strains were within the same branch; and the DBA1J and DBA2J strains formed their own grouping. However, there were some differences in the phylogenetic trees produced using the different types of genetic variants. The SNP- and SV-based analyses grouped C57BL/6J, B10J, and B10.D2 mice together, which reflects their close genetic relationship. However, the TR-based analysis placed C57BL/6J in a separate branch, which may be due to the use of C57BL/6J as the reference sequence. Also, the phylogenetic trees had different clustering patterns for CE, TH, SMJ and a few other strains. These differences may be attributed to the different mutational mechanisms underlying the generation of SNP, SV, and TR alleles and the timing of their occurrence during the evolution of the different strains ^24,25^.

### TR alleles for two biomedical phenotypes

To determine if this murine TR database could be used to identify unknown genetic factors for biomedical traits (i.e., account for some missing heritability), we investigated whether strain-specific high impact TR alleles could provide genetic candidates for two strain-specific phenotypes that were both identified over 40 years ago. In 1981, PL/J mice were found to produce sperm with a high frequency (42%) of morphologic abnormalities, which include having an abnormally shaped head or completely lacking a head. PL/J sperm also have a high frequency of aneuploidy and abnormal spindle formation, and a reduced rate of crossing over. Analysis of PL/J intercross progeny indicated that the PL/J genetic factors causing these abnormalities are recessive and oligogenic ^26,27^, but none have yet been identified. We identified a large PL/J-unique TR allele in *Prdm9* that alters the amino acids at positions 664 to 847 of the PL/J protein, which contains six zinc finger C2H2-type domains that are critical for DNA-binding (**Figure 5A**). *Prdm9* encodes a zinc finger protein that binds to DNA at specific sites and trimethylates histone H3 at lysines 4 and 36 (H3K4me3 and H3K36me3) ^28^. During meiosis, Prdm9 determines the location of recombination hotspots, which control the sites for genetic recombination. It also determines where programmed DNA double strand breaks (DSBs) occur, which give rise to genetic exchange between chromosomes. In *Prdm9* knockout (KO) mice, meiotic cells make DSBs at residual H3K4me3 sites, but they are not repaired successfully; this causes them to undergo pachytene arrest and apoptosis, which results in a failure to produce sperm and eggs ^29,30^. Given the *Prdm9* KO-induced effects on sperm, the PL/J *Prdm9* TR allele is a likely genetic contributor to its abnormal sperm production.

**Figure 5.**
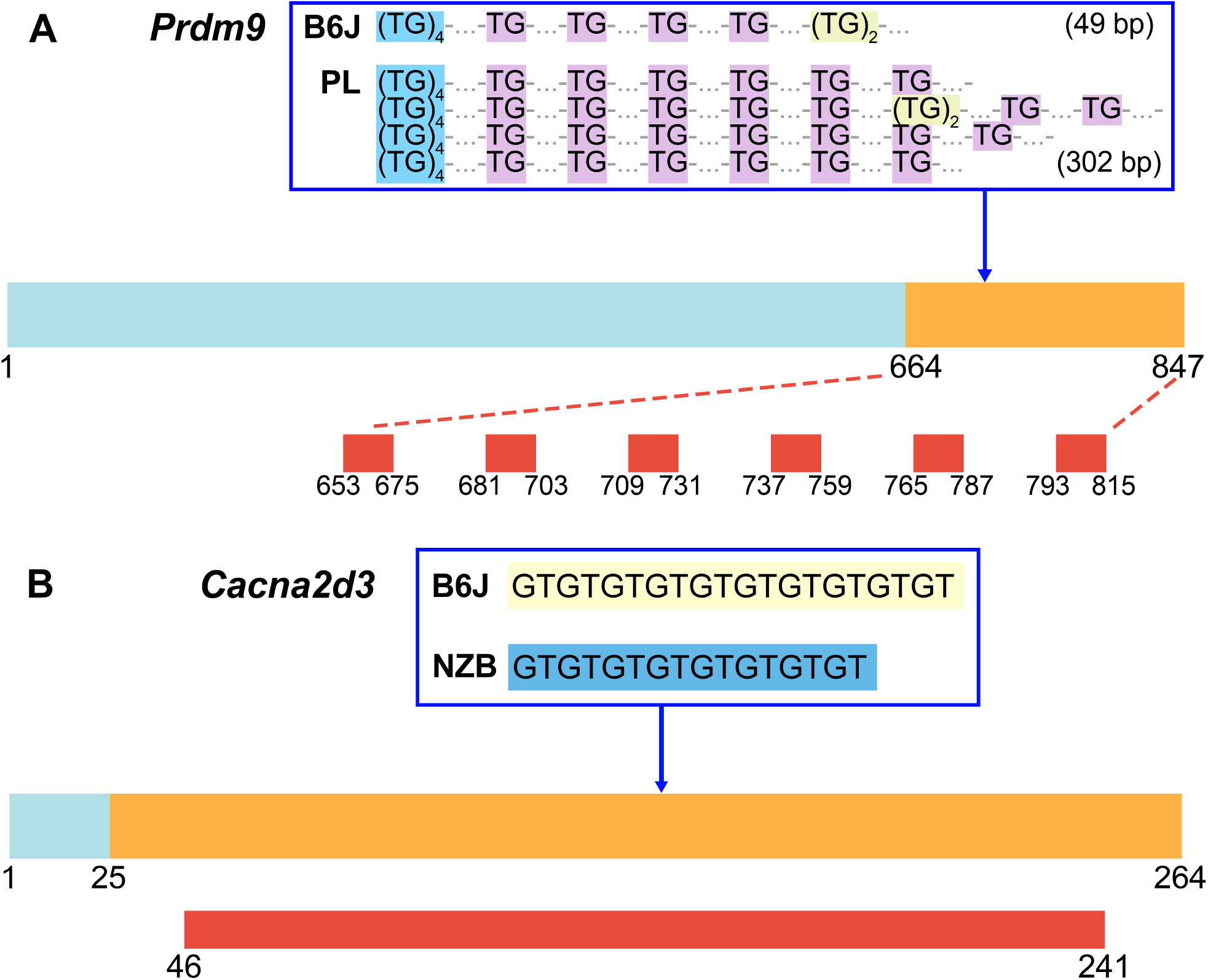
The effect of PL/J *Pdrm9* and NZB *Cacna2d3* TR alleles on protein structure. (A)The Prdm9 protein (residues 1-847) has a conserved *N*-terminal segment (1-664, blue), and a COOH terminal region (664-847, orange) whose sequence is altered by a PL/J-specific TR allele. The C57BL/6J (49 bp) and the PL/J (302 bp)TR alleles are shown above the protein diagram. The red rectangles (below) show the zinc finger C2H2-type domains in Prdm9 with the amino acid numbers for their starting and ending positions. The expanded PL/J TR allele alters the sequence of all six of these domains, which will greatly reduce Prdm9’s ability to bind to DNA. (B)The Cacna2d3 protein has a conserved *N*-terminal segment (residues 1-25, blue) and a COOH terminal region (residues 25-264, orange) whose sequence is altered by a NZB-specific TR allele. The C57BL/6 (GT)_10_ and NZB (GT)_8_ TR alleles are shown above the protein diagram, and a region with a conserved sequence is shown by the red rectangle below the protein. The NZB TR allele alters most of the amino acids in the Cacna2d3 protein sequence, which will compromise channel assembly and calcium conductance.

As a second example, NZB mice have developmental brain abnormalities that were noted in multiple papers published since 1985. The abnormalities consist of ectopic collections of layer 1 neurons with displacement of the underlying and adjacent cortical layers, which are often unilateral and located in somatosensory cortical areas ^31-33^. Moreover, NZB mice have a significant deficit in reversal learning, and exhibit a high level of spatial memory in the Morris water maze test ^34^. Although NZB mice are not a strain that is used for modeling Autism Spectral Disorder (ASD), their resistance to change a learned pattern of behavior reflects one feature of ASD. We identified a NZB-unique TR allele in *Cacna2d3*, which encodes the auxiliary (α2δ3) subunit of voltage-gated calcium channels (VGCCs) that are is expressed throughout the CNS ^35^. The NZB-unique TR allele alters amino acids 25 to 264, which is within a highly conserved region of Cacna2d3 (**Figure 5B**). Cacna2d3 regulates the surface expression and function of VGCCs, which is critical for neurotransmitter release; and it regulates synapse formation and synapse efficiency ^36^. *CACNA2D3* was identified as a potential cause of human ASD in multiple studies ^37-39^; and a conditional *Cacna2d3* knockout in parvalbumin-expressing interneurons produces key ASD behaviors that included an increase in repetitive behavior and improved spatial memory ^40^. Based upon the phenotypes exhibited by *Cacna2d3* knockout mice, the reversal learning deficits and high spatial memory exhibited by NZB mice are consistent with an effect of a NZB-unique TR allele that impairs Cacna2d3 function. This effect is also consistent with the recent finding that human TR alleles impact brain phenotypes, which include cortical surface area ^11^.

## DISCUSSION

Many properties of the murine TRs characterized here are consistent with those observed in humans and other species. (i) TRs are a source of genetic variation; they exhibit high rates of polymorphism within members of a species and are frequently multi-allelic. (ii) TRs result from chromosomal misalignment that leads to polymerase slippage, which generates stepwise changes in repeat numbers. (iii) TRs within protein coding regions or those that produce frameshift or termination mutations are rare ^41,42^. (iv) There is a high level of variation within TRs in the human genome, which is especially common in the non-coding regions of the genome ^43^. The mutation rate within human or yeast ^42^ TRs is 100 to 10,000-fold greater than that of SNPs, and TR mutations usually alter repeat copy number such that long alleles tend to contract and short alleles expand ^44^. Of note, the frequency of polymorphisms within TRs tend to correlate with paternal age ^45^,^44^. The high rate of polymorphism within TRs explains the relatively high number of alleles (n=6.52) per TR. The very low level of LD between murine TRs could be explained by the high level of polymorphism at TR sites that would disrupt LD between TRs.

We found that only 3.1% of murine TRs reside within protein-coding exons, whereas the majority are in intergenic (57%) or intronic regions (39%). This pattern suggests that the impact of murine TR alleles is not primarily to alter protein sequences, but they may play a role in modulating gene expression or chromatin structure ^46^. The motif length for 91% of the murine TRs contains four or fewer base pairs. Hence, a selection pressure favoring shorter motifs may be driven by a need to maintain replication fidelity since shorter motifs are less susceptible to replication slippage and introduction of structural variants, which minimizes the risk of mutational disruptions and enhances genome maintenance. In humans, only TRs with highly expanded repeats are linked with pathogenic outcomes, whereas many human repeat expansions do not cause disease ^10^. Our ability to characterize the impact of mouse (or other species) TR alleles is limited by the absence of computational tools for predicting their functional impact on a genome-wide scale. Machine learning algorithms ^47^ that were developed using human pathogenic loci as the training data set cannot be readily applied to murine datasets. The lack of positional correlation between murine TRs and LINE-1 elements indicates that we do not fully understand genomic context for TRs. However, murine TRs should be viewed as integral components of the genomic landscape that contribute to genetic diversity and evolutionary adaptability. Our analysis of two biomedical phenotypes in mouse strains, which were identified over 40 years ago but their genetic basis had not been determined, demonstrates the importance of characterizing TR alleles among the inbred strains. The impact of both of the identified TR alleles on the strain-specific phenotypes were validated by the effects observed in previously generated gene knockout mice. This work lays the foundation for future studies that will uncover the molecular mechanisms by which TRs influence genome stability, evolution and phenotypes. These investigations are essential for advancing fundamental biological research and for translational medicine.

### Limitations of the study

Despite the comprehensive identification of tandem repeats (TRs) across 40 inbred mouse strains using high-fidelity (HiFi) long-read sequencing, several limitations should be noted. First, since this TR catalog was generated by a reference-sequence guided assembly method, highly divergent TRs that are absent from the reference sequence may have escaped detection or could not be resolved. Second, since most murine TRs are intronic or intergenic, we cannot reliably predict the functional impact of most of the TRs that we identified. Understanding their effects on gene expression or chromatin structure will require experimental testing of the functional effect of selected TRs.

## RESOURCE AVAILABILITY

### Lead contact

Requests for further information and resources should be directed to and will be fulfilled by the lead contact, Gary Peltz.

### Materials availability

This study did not generate new unique reagents.

### Data and code availability

- The catalog and database of tandem repeats have been deposited in Zenodo and are publicly available at https://zenodo.org/records/15313223.
- The long-read sequencing (LRS) data have been deposited at the NCBI BioProject database under accession number PRJNA1250604 and are publicly available as of the date of publication.
- All software and analytical methods used in this study are publicly available, as listed in the key resources table.
- Any additional information required to reanalyze the data reported in this paper is available from the lead contact upon request.

## ACKNOWLEDGMENTS

This work was supported by NIH awards (1R01DC021133 and 1 R24 OD035408) to G.P.; The funder had no role in the writing of this paper. We thank Dr. Laura Reinholdt (Jackson Labs) for supplying the DNA obtained from several of the inbred strains.

## AUTHOR CONTRIBUTIONS

Conceptualization, G.P.; methodology, W.R., W.L, Z.F., and G.P.; formal analysis, software, and validation, W.R., W.L., E.D.,B.W., Z.C.; visualization, W.R.; writing—original draft, W.R. and G.P.; writing—review & editing, W.R. and G.P.; funding acquisition, G.P.; supervision, G.P.

## SUPPLEMENTAL INFORMATION

**Document S1. Figures S1-S3 and Tables S1 and S2**

**Table S3. List of strain-unique TRs located in exons or at transcription start sites, related to Figure 1 and 3**. The gene, chromosome, starting and ending position, motifs contained in the TR, genomic annotation, strains with the variant allele, and its predicted biotype are shown.

**Table S4. List of 30 TRs that were randomly selected for experimental validation, related to Figure 1**. The chromosome, start and end positions, reference and alternative TR alleles, motifs contained within each TR, strains with the variant allele, and the forward and reverse primer sequences used for their amplification are shown.

**Table S5. List of the TRs present in murine homologues of human TR expansion disease genes, related to Figure 1**. The human gene, disorder, murine homologues gene, chromosome, starting and ending position, reference and alternative allele, genomic location, and strains with the variant allele are shown.

## STAR★METHODS

## KEY RESOURCES TABLE

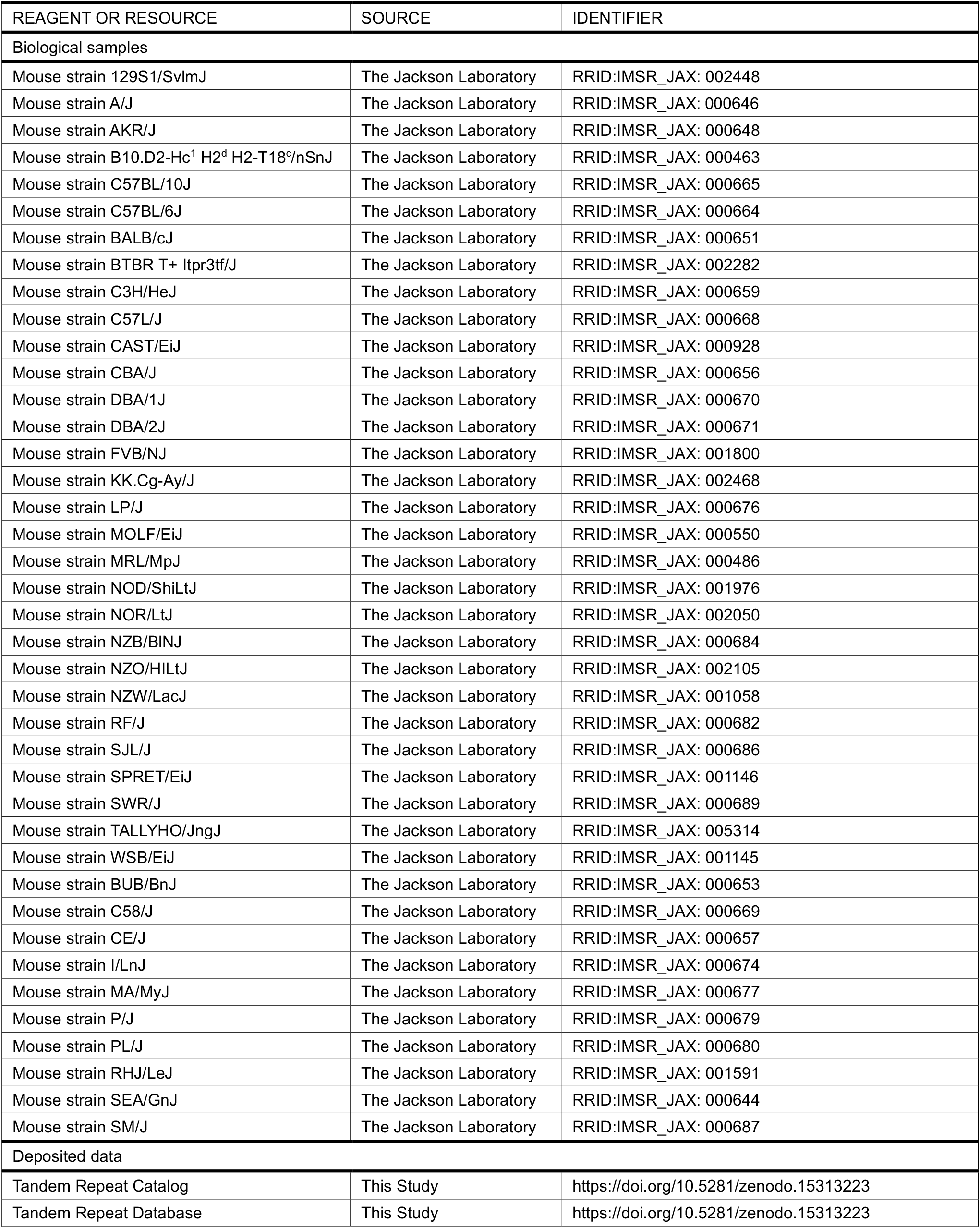
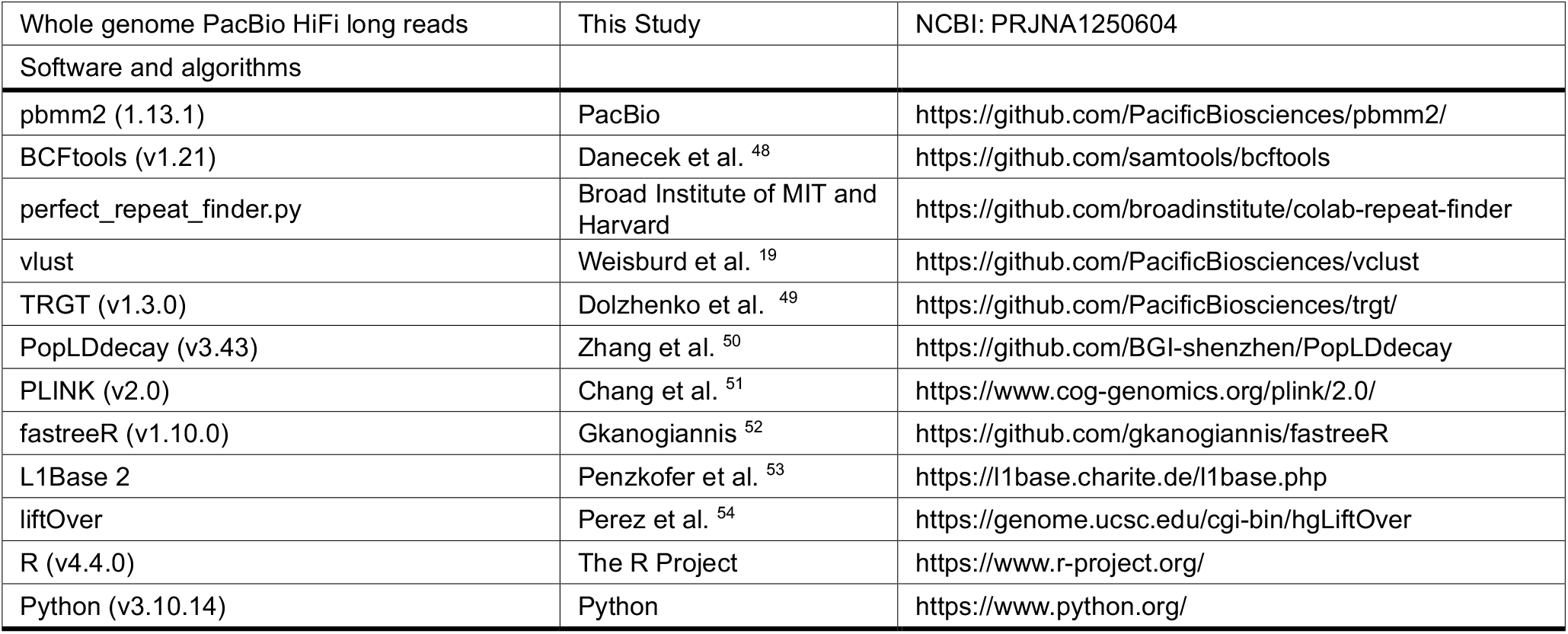

## METHOD DETAILS

### Animal experiments

All animal experiments were performed according to protocols that were approved by the Stanford Institutional Animal Care and Use Committee. All mice were obtained from Jackson Labs, and the results are reported according to the ARRIVE guidelines ^55^.

### Genomic Sequencing

Genomic DNA obtained from forty inbred strains (**Table S1**) were subject to LRS using the HiFi REVIO system (PacBIO) at the DNA Technologies Core of the Genome Center, University of California Davis, using methods that were fully described in ^18^.

### Generation of the TR catalog

Perfect repeats in the GRCm39 genome were identified using the Python script perfect_repeat_finder.py (https://github.com/broadinstitute/colab-repeat-finder) with the following parameters: a minimum repeat count of 3, a minimum spanning length of 9, a minimum motif size of 2, and a maximum motif size of 100. Subsequently, variation clusters--defined as contiguous regions containing variations across a given set of genome were detected using vlust ^19^. Finally, the output from the vlust command was converted into BED file format. Sample code is as follows:

python3 perfect_repeat_finder.py --min-repeats 3 --min-span 9 --min-motif-size 2 \

--max-motif-size 100 --output-prefix mm39.perfect.repeat \

--show-progress-bar mm39.fa

vclust --genome mm39.fa --reads strain1.bam strain2.bam strain3.bam … strain39.bam \

--regions mm39.perfect.repeat.bed > extended_regions txt

grep -v “NA” extended_regions.txt \

| awk ‘{OFS=“\t”; print $5, $1}’ | awk -F “[\t:-]” ‘{OFS=“\t”; print $1, $2, $3, $0}’ \

| cut -f 1-3,5 | sort -k 1,1 -k 2,2n -k 3,3n | bedtools merge -d -1 -c 4 -o distinct \

| awk ‘{OFS=“\t”; print $1, $2, $3, “ID=“$1”_”$2”_”$3”;MOTIFS=“$4”;STRUC=<TR>“}’ \ > trs.bed

### Genotype TR using TRGT

Using the TR catalog (named trs.bed), genotyping of the TR alleles in each of the 39 strains relative to the C57B/6 reference sequence was performed using TRGT (v1.3.0 ^19^. All single-sample VCFs were then merged into a joint multi-sample VCF using the ‘trgt merge’ command. Filtering was performed to exclude TRs that were non-polymorphic across all strains, i.e., those that shared the reference allele in every strain. TRs exhibiting a genotype pattern of n1/n2 (where n1 ≠ n2) in all strains, which was indicative of potentially mosaic TRs, were also excluded. The main example code is as follows:

trgt genotype --genome mm39.fa --repeats trs.bed \

--reads sample.align.sort.pbmm2.bam \

--output-prefix sample --threads 128

trgt merge --vcf *vcf.gz --genome mm39.fa --output-type z --output s39.vcf.gz

### Linkage disequilibrium (LD) decay

For analysis of the 35 inbred strains, PopLDdecay (v3.43) with default parameters were used to calculate LD decay for SNPs, SVs, and TRs ^50^. Four datasets were separately analyzed: 21 million SNPs, 220K SNPs, 220K SVs, and 1.8 million TRs. The 220K SNP subset was selected to assess how LD decay changes when the SNP density is reduced to levels comparable to that of SVs. The SV dataset included only deletions and insertions, and SV genotypes were treated as bi-allelic. Similarly, TR genotypes were also treated as bi-allelic although TRs exhibit greater allelic variation, for computational convenience alleles identical to the reference were coded as ‘0/0’, while those differing from the reference were coded as ‘1/1’. The LD decay plots were generated using the Plot_OnePop.pl script provided with PopLDdecay.

### Phylogenetic tree construction

For 40 mouse strains, we first performed LD pruning on the SNP, SV, and TR datasets separately using plink 2.0 with the parameters ‘--indep-pairwise 1000 100 0.2’, to enhance computational efficiency and more accurately reflect true evolutionary relationships^51^. Subsequently, a phylogenetic tree was constructed using the R package fastreeR (v1.10.0) ^52^.

### Distance calculation between TRs and LINE-1 elements

Mouse LINE-1 data were downloaded from L1Base 2^53^; the LINE-1 genome coordinates were converted from the GRCm38 to the GRCm39 reference genome^54^; and the positions of the TRs in our database were compared with those of the LINE-1 elements. Since the LINE-1 elements exceed 6000 bp in length, which is larger than nearly all the TRs, any TR located within a LINE-1 element was labelled as contained within that LINE-1. If either end of a LINE-1 element overlapped with a TR, it was classified as an overlap. Also, a TR was deemed proximal to a LINE-1 element if the distance from either end of the LINE-1 to the TR was <80 bp; whereas if the distance was >200bp, the TR was not labelled as proximal to the LINE-1.

### TR Validation

Genomic DNA was prepared from liver tissue obtained from AJ, B10J, CBA, NOD, TallyHO and C57BL/6J mice using PacBio’s Nanobind tissue kit according to the manufacturer’s instructions. For some strains, genomic DNA was prepared from tail tissue that was lysed in QuickExtract DNA Extraction Solution (Biosearch Technologies). PCR amplification of the sequences surrounding 30 selected TRs from genomic DNA was performed using the GoTaq G2 master mix (Promega) and the primers listed in **Table S4** according to the manufacturer’s instructions. Amplicons were separated and analyzed using agarose gels. PCR reactions were sent to McLab (South San Francisco, CA) for sanger sequencing. If the amplicons were >1 kb, additional internal primers were used for sequencing of those amplicons.

## REFERENCES

1 Tanudisastro, H. A., Deveson, I. W., Dashnow, H. & MacArthur, D. G. Sequencing and characterizing short tandem repeats in the human genome. Nat Rev Genet 25, 460–475 (2024). 10.1038/s41576-024-00692-3

2 Rajan-Babu, I. S., Dolzhenko, E., Eberle, M. A. & Friedman, J. M. Sequence composition changes in short tandem repeats: heterogeneity, detection, mechanisms and clinical implications. Nat Rev Genet 25, 476–499 (2024). 10.1038/s41576-024-00696-z

3 Hannan, A. J. Tandem repeat polymorphisms: modulators of disease susceptibility and candidates for ‘missing heritability’. Trends Genet 26, 59–65 (2010). 10.1016/j.tig.2009.11.008

4 Mukamel, R. E. et al. Protein-coding repeat polymorphisms strongly shape diverse human phenotypes. Science 373, 1499–1505 (2021). 10.1126/science.abg8289

5 Mukamel, R. E. et al. Repeat polymorphisms underlie top genetic risk loci for glaucoma and colorectal cancer. Cell 186, 3659–3673 e3623 (2023). 10.1016/j.cell.2023.07.002

6 English, A. C. et al. Analysis and benchmarking of small and large genomic variants across tandem repeats. Nat Biotechnol (2024). 10.1038/s41587-024-02225-z

7 Zhang, S. et al. Genome-wide investigation of VNTR motif polymorphisms in 8,222 genomes: Implications for biological regulation and human traits. Cell Genom, 100699 (2024). 10.1016/j.xgen.2024.100699

8 Paulson, H. Repeat expansion diseases. Handbook of clinical neurology 147, 105–123 (2018). 10.1016/B978-0-444-63233-3.00009-9

9 Zhou, Z. D., Jankovic, J., Ashizawa, T. & Tan, E. K. Neurodegenerative diseases associated with non-coding CGG tandem repeat expansions. Nat Rev Neurol 18, 145–157 (2022). 10.1038/s41582-021-00612-7

10 Depienne, C. & Mandel, J. L. 30 years of repeat expansion disorders: What have we learned and what are the remaining challenges? Am J Hum Genet 108, 764–785 (2021). 10.1016/j.ajhg.2021.03.011

11 Cui, Y. et al. Multi-omic quantitative trait loci link tandem repeat size variation to gene regulation in human brain. Nat Genet (2025). 10.1038/s41588-024-02057-2

12 Horton, C. A. et al. Short tandem repeats bind transcription factors to tune eukaryotic gene expression. Science 381, eadd1250 (2023). 10.1126/science.add1250

13 Zu, T. et al. Non-ATG-initiated translation directed by microsatellite expansions. Proc Natl Acad Sci U S A 108, 260–265 (2011). 10.1073/pnas.1013343108

14 Cleary, J. D., Pattamatta, A. & Ranum, L. P. W. Repeat-associated non-ATG (RAN) translation. J Biol Chem 293, 16127–16141 (2018). 10.1074/jbc.R118.003237

15 Zu, T. et al. Metformin inhibits RAN translation through PKR pathway and mitigates disease in C9orf72 ALS/FTD mice. Proc Natl Acad Sci U S A 117, 18591–18599 (2020). 10.1073/pnas.2005748117

16 Fang, Z. & Peltz, G. Twenty-first century mouse genetics is again at an inflection point. Lab Animal (2025). 10.1038/s41684-024-01491-3

17 Ren, W. et al. A Murine Database of Structural Variants Enables the Genetic Architecture of a Spontaneous Murine Lymphoma to be Characterized. BioRxiv https://biorxiv.org/cgi/content/short/2025.01.09.632219v1 (2025). https://doi.org:biorxiv.org/cgi/content/short/2025.01.09.632219v1

18 Ren, W. et al. A Murine Database of Structural Variants Enables the Genetic Architecture of a Spontaneous Murine Lymphoma to be Characterized. bioRxiv (2025). 10.1101/2025.01.09.632219

19 Weisburd, B. et al. Defining a tandem repeat catalog and variation clusters for genome-wide analyses and population databases. bioRxiv, 2024.2010.2004.615514 (2024). 10.1101/2024.10.04.615514

20 Weisburd, B. et al. Defining a tandem repeat catalog and variation clusters for genome-wide analyses and population databases. BioRxiv 10.04.615514 (2024). 10.1101/2024.10.04.615514

21 Ahmed, M. & Liang, P. Transposable elements are a significant contributor to tandem repeats in the human genome. Comp Funct Genomics 2012, 947089 (2012). 10.1155/2012/947089

22 Steely, C. J., Watkins, W. S., Baird, L. & Jorde, L. B. The mutational dynamics of short tandem repeats in large, multigenerational families. Genome Biol 23, 253 (2022). 10.1186/s13059-022-02818-4

23 Richardson, S. R. et al. The Influence of LINE-1 and SINE Retrotransposons on Mammalian Genomes. Microbiol Spectr 3, MDNA3–0061-2014 (2015). 10.1128/microbiolspec.MDNA3-0061-2014

24 Mortazavi, M. et al. SNPs, short tandem repeats, and structural variants are responsible for differential gene expression across C57BL/6 and C57BL/10 substrains. Cell Genom 2 (2022). 10.1016/j.xgen.2022.100102

25 Carvalho, C. M. & Lupski, J. R. Mechanisms underlying structural variant formation in genomic disorders. Nat Rev Genet 17, 224–238 (2016). 10.1038/nrg.2015.25

26 Burkhart, J. G. & Malling, H. V. Sperm Abnormalities in the PL/J Mouse Strain: A Desription and Proposed Mechanism for Malformation. Gamete Research 4, 171–183 (1981).

27 Pyle, A. & Handel, M. A. Meiosis in male PL/J mice: a genetic model for gametic aneuploidy. Mol Reprod Dev 64, 471–481 (2003). 10.1002/mrd.10231

28 Paigen, K. & Petkov, P. M. PRDM9 and Its Role in Genetic Recombination. Trends Genet 34, 291–300 (2018). 10.1016/j.tig.2017.12.017

29 Brick, K., Smagulova, F., Khil, P., Camerini-Otero, R. D. & Petukhova, G. V. Genetic recombination is directed away from functional genomic elements in mice. Nature 485, 642–645 (2012). 10.1038/nature11089

30 Thibault-Sennett, S. et al. Interrogating the Functions of PRDM9 Domains in Meiosis. Genetics 209, 475–487 (2018). 10.1534/genetics.118.300565

31 Sherman, G. F., Galaburda, A. M., Behan, P. O. & Rosen, G. D. Neuroanatomical anomalies in autoimmune mice. Acta Neuropathol 74, 239–242 (1987). 10.1007/BF00688187

32 Sherman, G. F., Galaburda, A. M. & Geschwind, N. Cortical anomalies in brains of New Zealand mice: a neuropathologic model of dyslexia? Proc Natl Acad Sci U S A 82, 8072–8074 (1985). 10.1073/pnas.82.23.8072

33 Zilles, K. in Senile dementia of the Alzheimer type (eds J. Traber & W.W. Gispen) 355–365 (Springer, 1985).

34 Moy, S. S. et al. Social approach and repetitive behavior in eleven inbred mouse strains. Behav Brain Res 191, 118–129 (2008). 10.1016/j.bbr.2008.03.015

35 Dolphin, A. C. Calcium channel auxiliary alpha2delta and beta subunits: trafficking and one step beyond. Nat Rev Neurosci 13, 542–555 (2012). 10.1038/nrn3311

36 Hoppa, M. B., Lana, B., Margas, W., Dolphin, A. C. & Ryan, T. A. alpha2delta expression sets presynaptic calcium channel abundance and release probability. Nature 486, 122–125 (2012). 10.1038/nature11033

37 De Rubeis, S. et al. Synaptic, transcriptional and chromatin genes disrupted in autism. Nature 515, 209–215 (2014). 10.1038/nature13772

38 Satterstrom, F. K. et al. Large-Scale Exome Sequencing Study Implicates Both Developmental and Functional Changes in the Neurobiology of Autism. Cell 180, 568–584 e523 (2020). 10.1016/j.cell.2019.12.036

39 Girirajan, S. et al. Refinement and discovery of new hotspots of copy-number variation associated with autism spectrum disorder. Am J Hum Genet 92, 221–237 (2013). 10.1016/j.ajhg.2012.12.016

40 Shao, W. et al. Deletions of Cacna2d3 in parvalbumin-expressing neurons leads to autistic-like phenotypes in mice. Neurochem Int 169, 105569 (2023). 10.1016/j.neuint.2023.105569

41 Verbiest, M. et al. Mutation and selection processes regulating short tandem repeats give rise to genetic and phenotypic diversity across species. J Evol Biol 36, 321–336 (2023). 10.1111/jeb.14106

42 Verstrepen, K. J., Jansen, A., Lewitter, F. & Fink, G. R. Intragenic tandem repeats generate functional variability. Nat Genet 37, 986–990 (2005). 10.1038/ng1618

43 Duitama, J. et al. Large-scale analysis of tandem repeat variability in the human genome. Nucleic Acids Res 42, 5728–5741 (2014). 10.1093/nar/gku212

44 Sun, J. X. et al. A direct characterization of human mutation based on microsatellites. Nat Genet 44, 1161–1165 (2012). 10.1038/ng.2398

45 Mitra, I. et al. Patterns of de novo tandem repeat mutations and their role in autism. Nature 589, 246–250 (2021). 10.1038/s41586-020-03078-7

46 Hannan, A. J. Tandem repeats mediating genetic plasticity in health and disease. Nat Rev Genet 19, 286–298 (2018). 10.1038/nrg.2017.115

47 Fazal, S. et al. RExPRT: a machine learning tool to predict pathogenicity of tandem repeat loci. Genome Biol 25, 39 (2024). 10.1186/s13059-024-03171-4

48 Danecek, P. et al. Twelve years of SAMtools and BCFtools. Gigascience 10 (2021). 10.1093/gigascience/giab008

49 Dolzhenko, E. et al. Characterization and visualization of tandem repeats at genome scale. Nat Biotechnol (2024). 10.1038/s41587-023-02057-3

50 Zhang, C., Dong, S. S., Xu, J. Y., He, W. M. & Yang, T. L. PopLDdecay: a fast and effective tool for linkage disequilibrium decay analysis based on variant call format files. Bioinformatics 35, 1786–1788 (2019). 10.1093/bioinformatics/bty875

51 Chang, C. C. et al. Second-generation PLINK: rising to the challenge of larger and richer datasets. Gigascience 4, 7 (2015). 10.1186/s13742-015-0047-8

52 Gkanogiannis, A. fastreeR: Phylogenetic, Distance and Other Calculations on VCF and Fasta Files. https://bioconductor.org/packages/fastreeR. R package version 1.12.0 (2025). 10.18129/B9.bioc.fastreeR

53 Penzkofer, T. et al. L1Base 2: more retrotransposition-active LINE-1s, more mammalian genomes. Nucleic Acids Res 45, D68–D73 (2017). 10.1093/nar/gkw925

54 Perez, G. et al. The UCSC Genome Browser database: 2025 update. Nucleic Acids Res 53, D1243–D1249 (2025). 10.1093/nar/gkae974

55 Kilkenny, C., Browne, W. J., Cuthill, I. C., Emerson, M. & Altman, D. G. Improving bioscience research reporting: the ARRIVE guidelines for reporting animal research. PLoS Biol 8, e1000412 (2010). 10.1371/journal.pbio.1000412

