## Supplemental information for "A Tandem Repeat Atlas for the Genome of Inbred Mouse Strains: A Genetic Variation Resource"

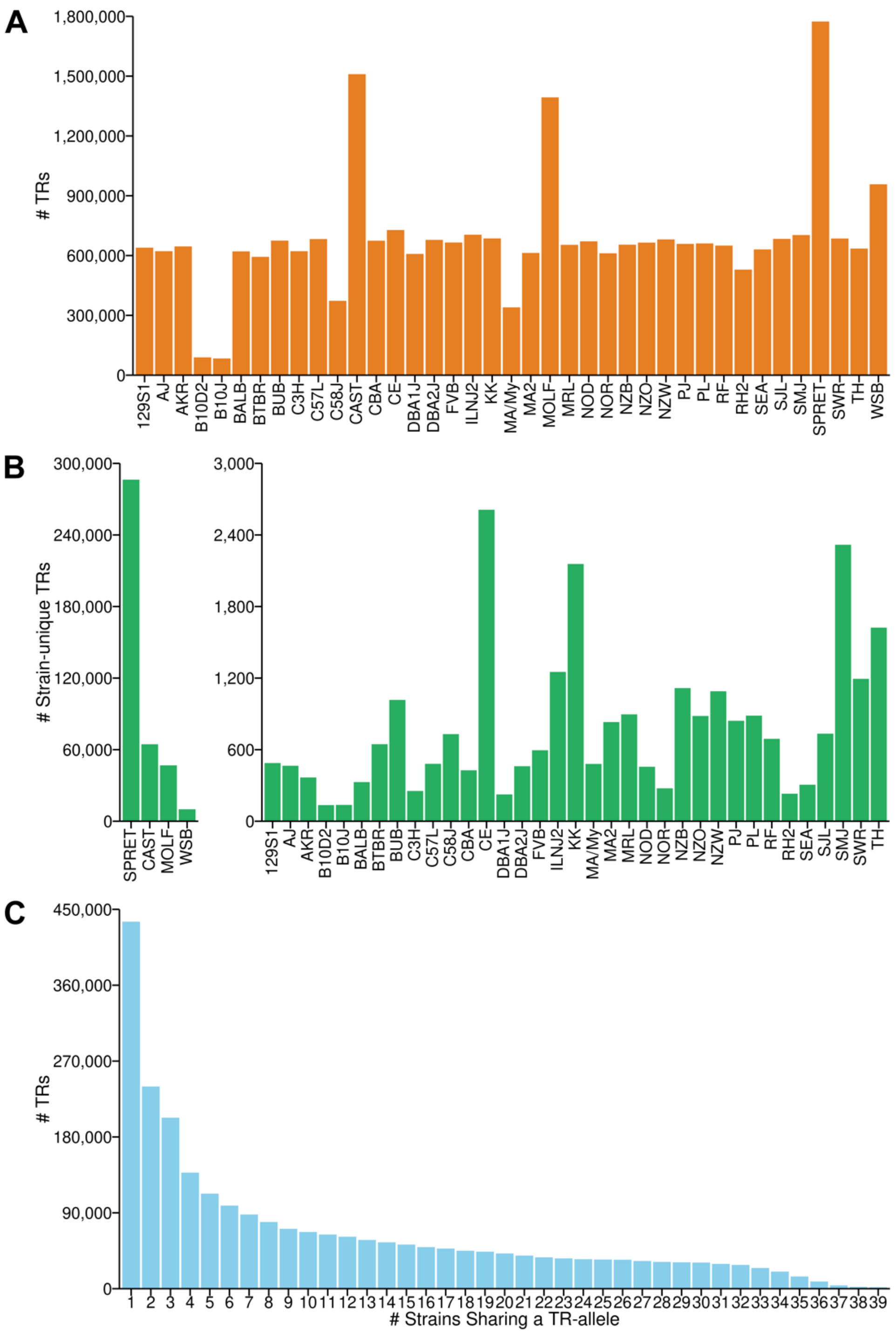


**Figure S1. Distribution and characteristics of TRs in all 39 strains, Related to Figures 1 and 2.** The total number of TRs (**A**) and the number of strain-unique TRs (**B**) are shown for each strain. The number of strain-unique TRs in the three wild-derived strains (SPRET, CAST and MOLF) (left) is significantly higher than in the other classical inbred strains (right). (**C**) The number of TRs where a minor allele is shared by the indicated number of strains is shown. Most of the minor TR alleles are shared by 1-3 strains.

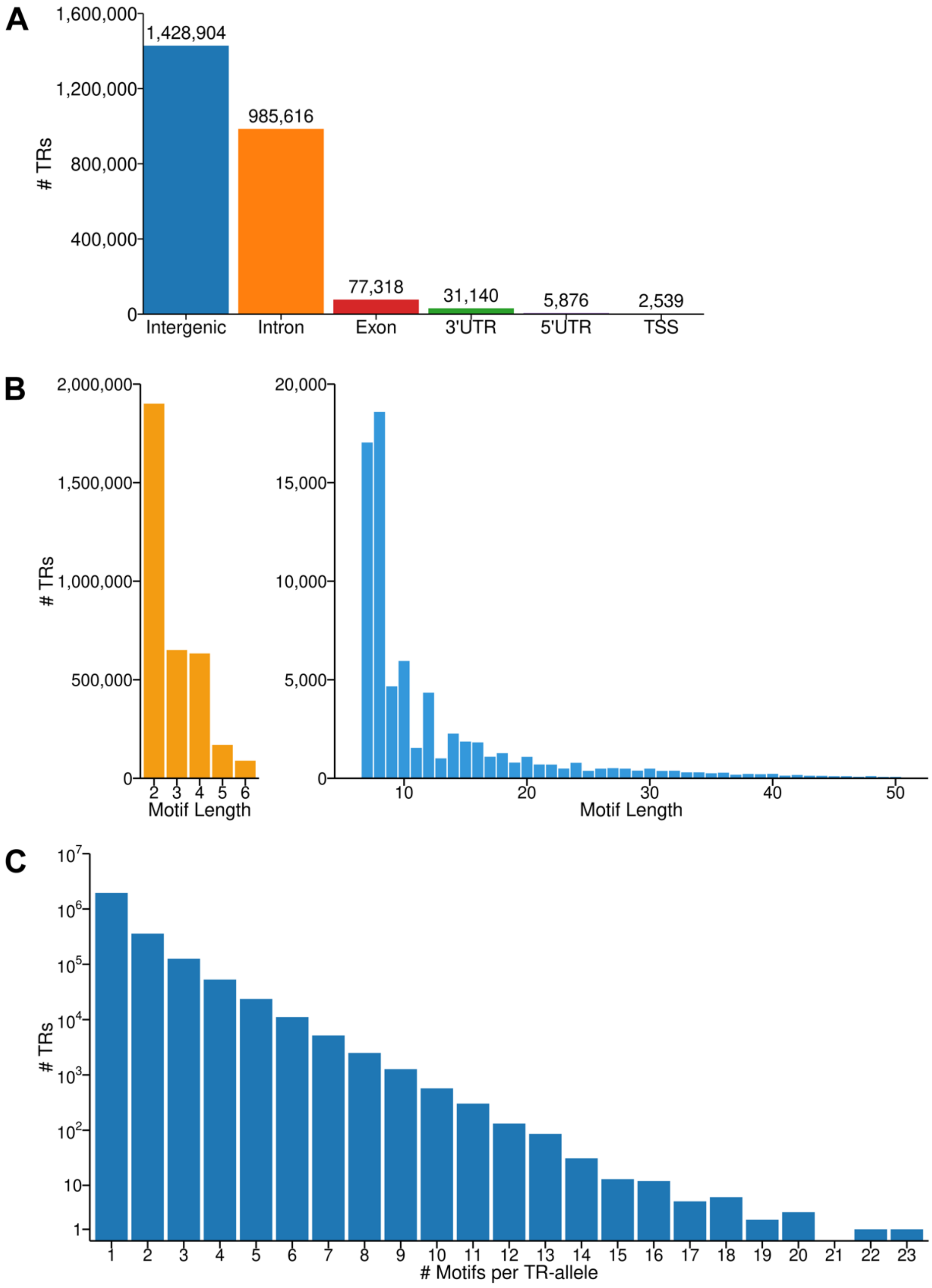


**Figure S2. The genomic distribution and properties of TRs in all 39 inbred strains, Related to Figures 1 and 3.** (**A**) The distribution of TRs in different types of genomic regions. (**B**) The number of TRs with different motif lengths. Most TRs are <7 bp (left), while TRs with motifs >6 bp are rarer (right). (**C**) The number of TRs with alleles with the indicated number of motifs. The Y-axis is $log_{10}$ transformed.

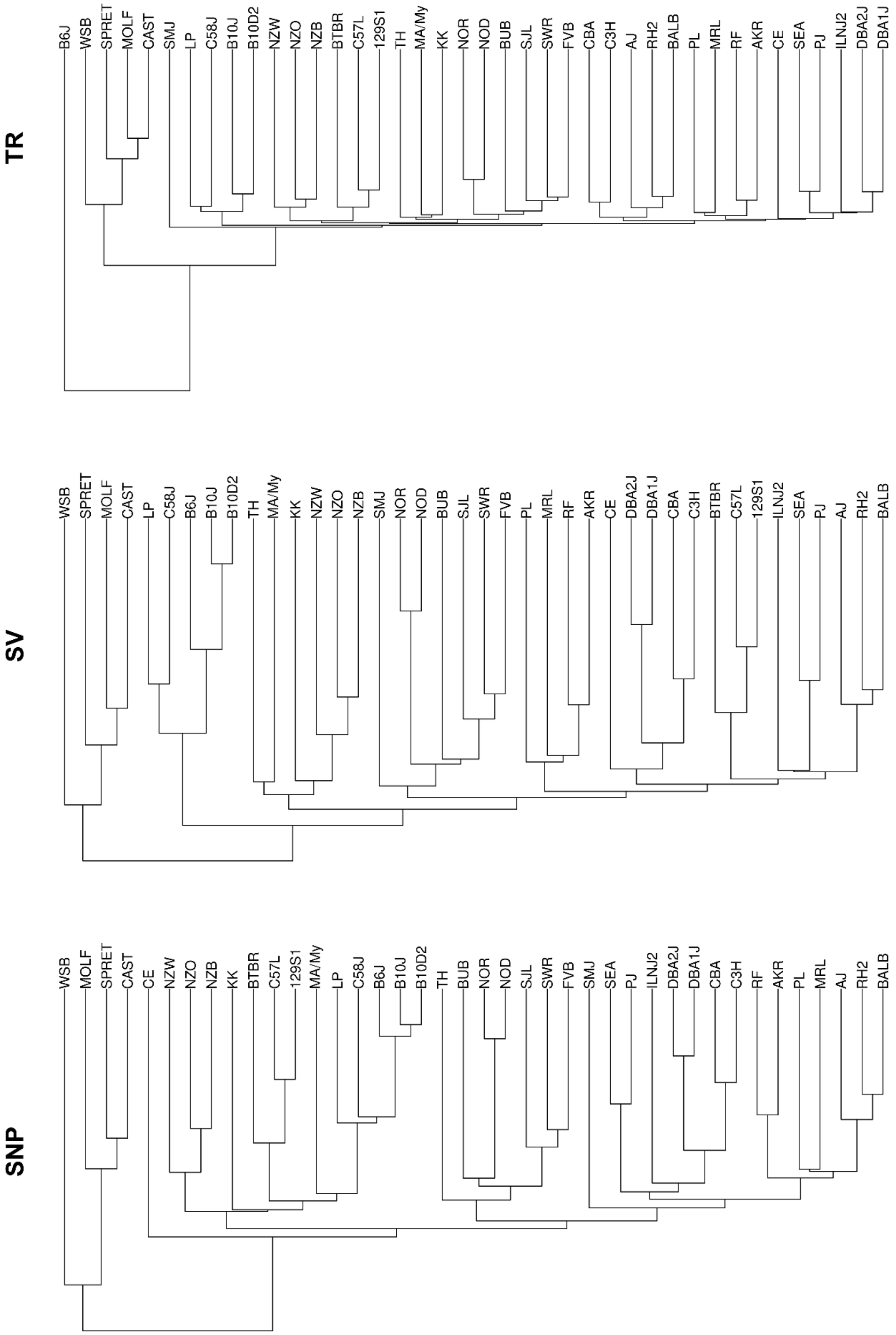


**Figure S3. Phylogenetic trees for 40 inbred mouse strains were constructed using SNP (n=294,695), structural variant (SV) (n=6,318), and TR (n=55,400) alleles. Related to Figures 1 and 4.**

**Table S1. Characteristics of the LRS data obtained from 39 inbred strains, Related to Figure 1.** The strain name, Jackson Lab #, sequence amount of sequence in gigabases (GB); and the # of reads (in millions) and mean read length mean read length are shown for each strain are shown. The number of TRs identified for each strain (relative to the C57BL/6 reference sequence) are also shown.

| Strain | JAX Strain # | GB | # Reads (M) | Mean Read Length | # TR |
|---|---|---|---|---|---|
| 129S1/SvlmJ | #002448 | 84.49 | 5.16 | 16387 | 639430 |
| A/J | #000646 | 85.12 | 4.69 | 18132 | 621763 |
| AKR/J | #000648 | 94.11 | 5.63 | 16725 | 645854 |
| B10.D2-Hc1 H2d H2-T18c/nSnJ | #000463 | 98.45 | 5.89 | 16722 | 89490 |
| C57BL/6J | #000664 | 87.65 | 5.05 | 17359 | Ref |
| C57BL/10J | #000665 | 92.54 | 5.76 | 16071 | 84069 |
| BALB/cJ | #000651 | 108.88 | 6.67 | 16333 | 620684 |
| BTBR T+ Itpr3tf/J | #002282 | 82.31 | 4.72 | 17433 | 593558 |
| BUB/BnJ | #000653 | 81.32 | 6.38 | 12741 | 675107 |
| C3H/HeJ | #000659 | 87.20 | 5.53 | 15754 | 621908 |
| C57L/J | #000668 | 92.20 | 5.92 | 15583 | 682717 |
| C58/J | #000669 | 79.26 | 7.21 | 10997 | 373240 |
| CAST/EiJ | #000928 | 79.22 | 5.94 | 13344 | 1509705 |
| CBA/J | #000656 | 88.19 | 5.12 | 17230 | 674328 |
| CE/J | #000657 | 74.83 | 6.10 | 12266 | 728155 |
| DBA/1J | #000670 | 109.58 | 6.58 | 16644 | 608060 |
| DBA/2J | #000671 | 106.62 | 6.03 | 17694 | 678487 |
| FVB/NJ | #001800 | 97.39 | 6.07 | 16039 | 665266 |
| I/LnJ | #000674 | 91.93 | 6.65 | 13829 | 704390 |
| KK.Cg-Ay/J | #002468 | 88.64 | 5.40 | 16426 | 685941 |
| LP/J | #000676 | 89.05 | 5.49 | 16210 | 340447 |
| MA/MyJ | #000677 | 87.84 | 6.22 | 14119 | 613075 |
| MOLF/EiJ | #000550 | 84.45 | 5.50 | 15345 | 1393391 |
| MRL/MpJ | #000486 | 96.00 | 6.17 | 15566 | 653598 |
| NOD/ShiLtJ | #001976 | 87.21 | 4.92 | 17720 | 671168 |
| NOR/LtJ | #002050 | 93.00 | 5.77 | 16114 | 611439 |
| NZB/BlNJ | #000684 | 87.23 | 5.27 | 16550 | 654656 |
| NZO/HILtJ | #002105 | 88.20 | 5.03 | 17538 | 665073 |
| NZW/LacJ | #001058 | 90.83 | 6.10 | 14884 | 680854 |
| P/J | #000679 | 92.88 | 5.80 | 16150 | 658581 |
| PL/J | #000680 | 70.32 | 5.96 | 11803 | 660916 |
| RF/J | #000682 | 92.80 | 5.49 | 16893 | 649910 |
| RHJ/LeJ | #001591 | 81.86 | 7.67 | 10668 | 529331 |
| SEA/GnJ | #000644 | 90.93 | 7.14 | 12737 | 630885 |
| SJL/J | #000686 | 101.17 | 6.45 | 15679 | 683630 |
| SM/J | #000687 | 46.31 | 4.70 | 9864 | 702692 |
| SPRET/EiJ | #001146 | 82.31 | 4.86 | 16936 | 1773873 |
| SWR/J | #000689 | 85.84 | 7.04 | 12194 | 685359 |
| TALLYHO/JngJ | #005314 | 85.01 | 5.59 | 15211 | 635120 |
| WSB/EiJ | #001145 | 97.26 | 6.03 | 16135 | 957544 |
| **Total** | | **3540.45** | **233.69** | **15301** | |
| **Mean** | | **88.51** | **5.84** | | |

**Table S2. The number of different categories of TRs identified in each of the 39 strains, Related to Figure 1.** The number of homozygous alternative alleles (Homo ALT), those matching the C57BL/6 sequence (Homo Ref) and TRs that could not be assessed (NA) by the TRGT program are indicated for each strain. Also shown are the number of TRs with heterozygous genotypes (i.e., where one genotype matched (0/1) or did not match (1/2) the TR in the reference strain).

| Strain | Homo ALT | Homo Ref | Het Alt (0/1) | Het Alt (1/2) | NA |
|---|---|---|---|---|---|
| 129S1 | 639430 | 2484773 | 150020 | 203647 | 17031 |
| AJ | 621763 | 2470928 | 167811 | 218489 | 15910 |
| AKR | 645854 | 2510011 | 133753 | 190984 | 14299 |
| B10D2 | 89490 | 3069753 | 264245 | 70207 | 1206 |
| B10J | 84069 | 3079079 | 261108 | 67833 | 2812 |
| BALB | 620684 | 2553218 | 129641 | 174909 | 16449 |
| BTBR | 593558 | 2594504 | 133902 | 158295 | 14642 |
| BUB | 675107 | 2505806 | 113586 | 184556 | 15846 |
| C3H | 621908 | 2481805 | 160024 | 216616 | 14548 |
| C57L | 682717 | 2495626 | 120644 | 178419 | 17495 |
| C58J | 373240 | 2786903 | 189054 | 136235 | 9469 |
| CAST | 1509705 | 1547742 | 24838 | 290348 | 122268 |
| CBA | 674328 | 2495122 | 125637 | 183028 | 16786 |
| CE | 728155 | 2420209 | 111563 | 215381 | 19593 |
| DBA1J | 608060 | 2487564 | 167184 | 215509 | 16584 |
| DBA2J | 678487 | 2534899 | 107273 | 156113 | 18129 |
| FVB | 665266 | 2519770 | 115011 | 178657 | 16197 |
| ILNJ2 | 704390 | 2494259 | 103685 | 172614 | 19953 |
| KK | 685941 | 2475677 | 121436 | 191984 | 19863 |
| LP | 340447 | 2796721 | 215075 | 133370 | 9288 |
| MA/My | 613075 | 2545332 | 136847 | 182668 | 16979 |
| MOLF | 1393391 | 1624770 | 46222 | 315293 | 115225 |
| MRL | 653598 | 2491334 | 130533 | 200702 | 18734 |
| NOD | 671168 | 2499393 | 126234 | 178602 | 19504 |
| NOR | 611439 | 2562875 | 134340 | 168735 | 17512 |
| NZB | 654656 | 2457113 | 151399 | 213091 | 18642 |
| NZO | 665073 | 2501574 | 130362 | 180750 | 17142 |
| NZW | 680854 | 2473372 | 129156 | 191489 | 20030 |
| PJ | 658581 | 2534691 | 117833 | 167161 | 16635 |
| PL | 660916 | 2488181 | 125108 | 201545 | 19151 |
| RF | 649910 | 2486733 | 144803 | 197213 | 16242 |
| RH2 | 529331 | 2621757 | 158133 | 172637 | 13043 |
| SEA | 630885 | 2550222 | 118306 | 180600 | 14888 |
| SJL | 683630 | 2519814 | 104225 | 168852 | 18380 |
| SMJ | 702692 | 2467526 | 110575 | 191805 | 22303 |
| SPRET | 1773873 | 1054130 | 28303 | 349895 | 288700 |
| SWR | 685359 | 2485722 | 112235 | 193690 | 17895 |
| TH | 635120 | 2479378 | 152090 | 209601 | 18712 |
| WSB | 957544 | 2196506 | 60037 | 246759 | 34055 |

**Table S3. List of strain-unique TRs located in exons or at transcription start sites, related to Figure 1 and 3.**

| Gene | Chrom | Start | End | Motifs | Annotation | Strain | Biotype |
|---|---|---|---|---|---|---|---|
| Aff3 | 1 | 38257463 | 38257487 | GCC,GCT | exon | SWR | protein coding gene |
| Cdc73 | 1 | 143578466 | 143578488 | CCG | exon | MRL | protein coding gene |
| Catspere2 | 1 | 177819065 | 177819446 | CC,GAG,TA,TGC | exon | PJ | protein coding gene |
| Cenpf | 1 | 189417141 | 189417160 | CTTCTG | exon | ILNJ2 | protein coding gene |
| Armt1 | 10 | 4382567 | 4382592 | AA | exon | NOD | protein coding gene |
| Plagl1 | 10 | 13003773 | 13003817 | GCCAATGCA | exon | BUB | protein coding gene |
| Plagl1 | 10 | 13003836 | 13003871 | GCCAATGCA | exon | BUB | protein coding gene |
| Raet1e | 10 | 22057965 | 22057984 | ATTTGC | exon | KK | protein coding gene |
| Sowahc | 10 | 59059313 | 59059340 | GGA | exon | SMJ | protein coding gene |
| Cactin | 10 | 81157152 | 81157207 | CGGAGC,CGGAGT | exon | BUB | protein coding gene |
| Ptprb | 10 | 116119582 | 116119662 | AGACCCTCGGGAGCACTGCAA | exon | SMJ | protein coding gene |
| Purb | 11 | 6425830 | 6425850 | CCG | exon | KK | protein coding gene |
| Zcchc10 | 11 | 53223344 | 53223365 | CAG | exon | KK | protein coding gene |
| Or1e30 | 11 | 73676994 | 73678077 | TA,TCA | exon | CE | protein coding gene |
| Cpd | 11 | 76737703 | 76737724 | AGC | exon | NZW | protein coding gene |
| Nufip2 | 11 | 77577146 | 77577168 | ATT,CAC | exon | BUB | protein coding gene |
| Nufip2 | 11 | 77577186 | 77577205 | ACC | exon | BUB | protein coding gene |
| Supt6 | 11 | 78123569 | 78123588 | CTT,TCA | exon | NZW | protein coding gene |
| Krt24 | 11 | 99175704 | 99175759 | CCCCCACCAAAGCCAGAG | exon | KK | protein coding gene |
| Cep95 | 11 | 106704660 | 106704685 | GAGGAA,GGA | exon | SJL | protein coding gene |
| Tex19.1 | 11 | 121037811 | 121037837 | GAGGAA | exon | SJL | protein coding gene |
| Ccdc177 | 12 | 80805733 | 80805756 | GAGGCT | exon | KK | protein coding gene |
| Rgs6 | 12 | 83184397 | 83184418 | AA | exon | SWR | protein coding gene |
| Nek9 | 12 | 85353076 | 85353109 | CCA,CCG | exon | CE | protein coding gene |
| Ak7 | 12 | 105676448 | 105676468 | GAG | exon | KK | protein coding gene |
| Ahnak2 | 12 | 112743219 | 112744264 | CAG | exon | TH | protein coding gene |
| Ahnak2 | 12 | 112747576 | 112747603 | CAG | exon | FVB | protein coding gene |
| Ahnak2 | 12 | 112748005 | 112748429 | CAG | exon | CE | protein coding gene |
| Ighv2-9-1 | 12 | 113732091 | 113734647 | TC | exon | AJ | gene segment |
| Ighv1-63 | 12 | 115456904 | 115459601 | GA | exon | CE | gene segment |
| Stard3nl | 13 | 19557325 | 19557521 | AC,AGT,AT,GA | exon | DBA2J | protein coding gene |
| H2ac10 | 13 | 23718423 | 23718514 | TGC,TT | exon | NOR | protein coding gene |
| Simc1 | 13 | 54672827 | 54672965 | CAGGAAGTGTGACCCAGTCAC TAAGAAGTGTGATGCAGTCAT | exon | BTBR | protein coding gene |
| Tut7 | 13 | 59947767 | 59947791 | TCA,TCC | exon | MRL | protein coding gene |
| Cacna2d3 | 14 | 28691248 | 28691268 | GT | exon | NZB | protein coding gene |
| Pabpn1 | 14 | 55131614 | 55131644 | GCA,GGC | exon | NZB | protein coding gene |
| Dach1 | 14 | 98406277 | 98406346 | CTG,GCT,GCTGCTGCTGCTAC TGCT | exon | SWR | protein coding gene |
| Col14a1 | 15 | 55249481 | 55249632 | AC,AG | exon | C57L | protein coding gene |
| Fam83h | 15 | 75874624 | 75874752 | CTCCCCTTGCGCTCAGGGTAA GCTGGGGTAGGG | exon | SJL | protein coding gene |
| Dgat1 | 15 | 76395814 | 76395834 | GGAGCC | exon | MA/My | protein coding gene |
| Cdc42ep1 | 15 | 78733421 | 78733442 | ACCGTG | exon | MA/My | protein coding gene |
| Kcnh3 | 15 | 99130669 | 99130692 | GCA | exon | SMJ | protein coding gene |
| Crebbp | 16 | 3902589 | 3902616 | TGC,TTG | exon | SMJ | protein coding gene |
| Pi4ka | 16 | 17223602 | 17223626 | GCC | exon | ILNJ2 | protein coding gene |
| Crygs | 16 | 22624324 | 22624357 | CAA | exon | TH | protein coding gene |
| Heg1 | 16 | 33505019 | 33505040 | GCT | exon | SMJ | protein coding gene |
| Muc13 | 16 | 33619370 | 33619812 | AGTCAATCTCCAGGTAGTTCA TCTCAGGCCTCTACTACAACAT CGTCTTCTGGTGGCGCCAGTC CTCCCACCACGGTACAG | exon | SMJ | protein coding gene |
| Zbtb21 | 16 | 97752887 | 97752911 | GCT | exon | NZO | protein coding gene |
| Arid1b | 17 | 5045567 | 5045587 | CAG,GCC | exon | 129S1 | protein coding gene |
| Tmem181a | 17 | 6347857 | 6349375 | CAGA | exon,TSS | PL | protein coding gene |
| Prdm9 | 17 | 15764834 | 15764882 | TG | exon | PL | protein coding gene |
| Brd4 | 17 | 32417124 | 32417147 | CTG | exon | PL | protein coding gene |
| Brd4 | 17 | 32417154 | 32417180 | CTG | exon | PL | protein coding gene |
| Esp31 | 17 | 38951138 | 38952048 | ATA | exon | ILNJ2 | protein coding gene |
| Esp31 | 17 | 38955373 | 38955663 | TT | exon | ILNJ2 | protein coding gene |
| Pcare | 17 | 72051705 | 72051728 | CTG | exon | SMJ | protein coding gene |
| Ecscr | 18 | 35849813 | 35849845 | CTG | exon | SMJ | protein coding gene |
| Prdm6 | 18 | 53597807 | 53597827 | CCG,CCT | exon | KK | protein coding gene |
| Cep192 | 18 | 67957837 | 67957873 | TGG | exon | SMJ | protein coding gene |
| Rbm4 | 19 | 4837606 | 4837633 | AGC,GCT | exon | LP | protein coding gene |

| | | | | | | | |
|---|---|---|---|---|---|---|---|
| Tm7sf2 | 19 | 6113092 | 6113191 | ATGGGATAGCCCTGGAGGGAAGGGACCC | exon | BUB | protein coding gene |
| Tm7sf2 | 19 | 6116142 | 6116165 | CT | exon | BUB | protein coding gene |
| Hnrnpul2 | 19 | 8797867 | 8797899 | GAC,GAG | exon | LP | protein coding gene |
| Gldc | 19 | 30152520 | 30152548 | CCG,CCGCCA | exon | MRL | protein coding gene |
| Itprip | 19 | 47886378 | 47886398 | CTC,GCT | exon | ILNJ2 | protein coding gene |
| Skida1 | 2 | 18052212 | 18052235 | GCA,GCC | exon | SJL | protein coding gene |
| Dnajc1 | 2 | 18221879 | 18221947 | TTCT,TTTC | three_prime_UTR,TSS | SJL | protein coding gene |
| Snapc4 | 2 | 26259538 | 26259557 | CTG | exon | MA/My | protein coding gene |
| Or1l4 | 2 | 37091265 | 37091285 | CAG | exon | TH | protein coding gene |
| Olfml2a | 2 | 38841373 | 38841409 | CAGACAGGAGGC | exon | BUB | protein coding gene |
| Olfml2a | 2 | 38844663 | 38844684 | CAC,CAG | exon | BUB | protein coding gene |
| Olfml2a | 2 | 38844689 | 38844736 | CAC,GCACCACCA | exon | BUB | protein coding gene |
| Slc43a3 | 2 | 84780349 | 84780378 | AGC | exon | FVB | protein coding gene |
| Phf21a | 2 | 92150728 | 92150758 | GCA | exon | AJ | protein coding gene |
| Shld1 | 2 | 132534028 | 132534272 | AA,CCCT,CTCCCC,CTCTCC,TC,TTC | exon | 129S1 | protein coding gene |
| Chgb | 2 | 132634838 | 132634935 | CAC,GAGGAAGGC | exon | ILNJ2 | protein coding gene |
| Mybl2 | 2 | 162916399 | 162916419 | AGG,GCA | exon | KK | protein coding gene |
| Svs4 | 2 | 164119092 | 164119121 | CTA,CTT | exon | KK | protein coding gene |
| Taf4 | 2 | 179617457 | 179617481 | CCG | exon | KK | protein coding gene |
| Kcnn3 | 3 | 89427972 | 89427994 | GCA | exon | KK | protein coding gene |
| Kcnn3 | 3 | 89427996 | 89428015 | GCA | exon | KK | protein coding gene |
| Flg2 | 3 | 93114122 | 93114160 | CGGTCA,GGTCAG | exon | KK | protein coding gene |
| Flg2 | 3 | 93114362 | 93114394 | CGGTCA,GGTCAG | exon | KK | protein coding gene |
| Flg2 | 3 | 93114596 | 93114628 | CGGTCA,GGTCAG | exon | ILNJ2 | protein coding gene |
| Flg2 | 3 | 93123531 | 93126825 | CGGTCA,GGTCAG | exon | BTBR | protein coding gene |
| Flg | 3 | 93188729 | 93191398 | GAG,GCA,GCACCA | exon | B10J | protein coding gene |
| BC028528 | 3 | 95795448 | 95795502 | TCACTGGTTCTGTGG | exon | TH | protein coding gene |
| Ubl4b | 3 | 107461752 | 107461775 | CCT | exon | CE | protein coding gene |
| Skint6 | 4 | 112668840 | 112672891 | AA,AAATA,AAC,CAA,CC | exon | C58J | protein coding gene |
| Skint6 | 4 | 112942864 | 112946880 | AC,TA | exon | NOD | protein coding gene |
| Skint6 | 4 | 113013093 | 113017153 | ATA,ATT,CC,GAA | exon | KK | protein coding gene |
| Skint5 | 4 | 113480373 | 113481259 | TT | exon | KK | protein coding gene |
| Skint5 | 4 | 113748026 | 113752711 | AA,AC,ATA,CT,TC,TT | exon | NZO | protein coding gene |
| Arid1a | 4 | 133479879 | 133479902 | GCC | exon | FVB | protein coding gene |
| Plekhg5 | 4 | 152197066 | 152197204 | AGG | exon | 129S1 | protein coding gene |
| Tnfrsf25 | 4 | 152201056 | 152201082 | CTG | exon | 129S1 | protein coding gene |
| Espn | 4 | 152209454 | 152209478 | TCC | exon | 129S1 | protein coding gene |
| Gatad1 | 5 | 3697436 | 3697469 | CCG,GGT | exon | CE | protein coding gene |
| Paxip1 | 5 | 27971044 | 27971084 | GCT,TGC | exon | BUB | protein coding gene |
| Epha5 | 5 | 84565201 | 84565230 | CGCT | exon | SMJ | protein coding gene |
| Rchy1 | 5 | 92110578 | 92110597 | CTC | exon,TSS | RF | protein coding gene |
| Vsig10 | 5 | 117489672 | 117489705 | GAA,GAG | exon | RF | protein coding gene |
| Slc8b1 | 5 | 120665699 | 120665855 | CC,GTG | exon | PL | protein coding gene |
| Zfp316 | 5 | 143250148 | 143250184 | CAC,CAT,TCA | exon | CBA | protein coding gene |
| Trrap | 5 | 144727801 | 144727822 | CCACCT | exon | TH | protein coding gene |
| Cdx2 | 5 | 147238838 | 147238858 | CTT,TGC | exon | TH | protein coding gene |
| Dlx6 | 6 | 6863493 | 6863513 | CAG | exon | PJ | protein coding gene |
| Nup50l | 6 | 96142059 | 96142106 | CCT,TCC | exon | SWR | protein coding gene |
| Peg3 | 7 | 6712167 | 6712265 | TGGGGCTCCTGGCCATGGGGCTTATCATCA | exon | SMJ | protein coding gene |
| Sult2a5 | 7 | 13395808 | 13396792 | AAC,AC,ACA,CTG,TC | exon | C58J | protein coding gene |
| Or51q1 | 7 | 103628624 | 103628715 | TTC | exon | TH | protein coding gene |
| Taok2 | 7 | 126473907 | 126473928 | TCC | exon | TH | protein coding gene |
| Srcap | 7 | 127157462 | 127157481 | TCC | exon | TH | protein coding gene |
| Or13a21 | 7 | 140001188 | 140001848 | AAGA,AAGG,AGAA,AGGG,GA | five_prime_UTR,TSS | BUB | protein coding gene |
| Nlrp6 | 7 | 140504026 | 140504047 | AGA | exon | NZW | protein coding gene |
| Deaf1 | 7 | 140907198 | 140907219 | GGC | exon | NZW | protein coding gene |
| Muc2 | 7 | 141292956 | 141293020 | CAA | exon | DBA1J | protein coding gene |
| Dusp8 | 7 | 141635788 | 141635927 | ACT,CCA,CTG,GCC,GCT,TGC | exon | NZW | protein coding gene |
| Irs2 | 8 | 11058346 | 11058367 | GTT | exon | TH | protein coding gene |
| Brf2 | 8 | 27614197 | 27614221 | CTG | exon | PL | protein coding gene |
| Cmtm1 | 8 | 105036334 | 105036487 | CCGGGTACTGAAGGTCCCTGGCTGGCTGGTGTC,CTGG | exon | PJ | protein coding gene |
| Terf2 | 8 | 107823023 | 107823050 | CGCCCTCCC | exon | RH2 | protein coding gene |
| Zfhx3 | 8 | 109674153 | 109674188 | CAG | exon | 129S1 | protein coding gene |
| Gse1 | 8 | 121297762 | 121297785 | CCA | exon | ILNJ2 | protein coding gene |
| Ctu2 | 8 | 123205723 | 123205749 | GCA | exon | ILNJ2 | protein coding gene |

| | | | | | | | |
|---|---|---|---|---|---|---|---|
| Ttc13 | 8 | 125448671 | 125448690 | GCAGCC | exon | NZB | protein coding gene |
| Rbm34 | 8 | 127696834 | 127696855 | TCA | exon | KK | protein coding gene |
| Ccdc7a | 8 | 129622575 | 129622597 | TTTC | exon | KK | protein coding gene |
| Eomes | 9 | 118307805 | 118307825 | GGA | exon | MA/My | protein coding gene |
| Kdm6a | X | 18029272 | 18029300 | CCG | exon | NZW | protein coding gene |
| Ids | X | 69408486 | 69408505 | GGC | exon | 129S1 | protein coding gene |
| Gabra3 | X | 71699742 | 71699901 | CCCT,CCTC,CT,GCTA | five_prime_UTR,TSS | NOR | protein coding gene |
| Dkc1 | X | 74139547 | 74139567 | TCC | five_prime_UTR,TSS | ILNJ2 | protein coding gene |
| Trap1a | X | 138237725 | 138237754 | GAA,GAG | exon | PL | protein coding gene |
| Ammecr1 | X | 141749439 | 141749474 | GCC | exon | KK | protein coding gene |
| Zrsr2 | X | 162719534 | 162719558 | CTGCGG | exon | TH | protein coding gene |

**Table S4. List of 30 TRs that were randomly selected for experimental validation, related to Figure 1.**

| Chrom | Start | End | Ref | Alt | Motifs | Strain | Forward Primer 1 Sequence | Reverse Primer 2 Sequence |
|---|---|---|---|---|---|---|---|---|
| chr1 | 119352843 | 119352905 | CTGTGTGTGTGTGTGTGTGTGTGTGTGTGTGTGTGTGTGTGTACATGTGTGTGTGTGTGTGTG | CTGTGTGTGTGTGTGTGTGTGTGTG | TG | AJ | CGTAAGGTTCTGGCTTGACACTATCA | CAAGCTCATGGATTCTGGAGGTC |
| chr2 | 174271744 | 174271769 | CCGAAAAAAAAAAAAAAAAAAAAAAA | CGGAAAAAAGAAAAAAAGAAA | AA | AJ | GATAGTCTATTCAGAAAGTTCCAGCCTATC | CTGACATAGTAACAGACAGATGGACAGA |
| chr11 | 25214764 | 25214812 | GGTGTGTGTGTGTGTGTGTGTGTGTGTGTGTGTGTGTGTGTGTGTGTGTGT | GGTGTTTGTGTGTGTGTGTGTGTGTGTGTGTGTGTGTATGTGT | GT | AJ | CAGGACATGAACTTCCTTCTGCTATT | GTGGCTCTATTAGGAAGCAGTGGA |
| chr1 | 76845972 | 76845998 | ACATGAGTGTGTGTGTGTGTGTGTGTG | ACATGAGTGTGTGTGTGTGTGTGTGTGTGTGTG | GT | AJ | GTCTGTTTTACAATCTAAAGGCACTCTG | GTCATTGTCCTTTATCCAAAAGTTAGAGAC |
| chr10 | 123414343 | 123414377 | CATTATTATTATTATTATTATTATTATTATTATTA | CACTATTATTATTATTATTATTATTA | ATT | AJ | GTTCTTATCCTTTGGCTTCTCCTAGG | ACACCATTGGAAAAGAGAACCTGG |
| chr6 | 29099453 | 29099501 | ATGTGTGTGTGTGTGTGTGTGTGTGTGTGTGTGTGTGTGTGTGTGTGTG | ATGTGTGTGTGTGTGTGTGTG | TG | AJ | CTTGGTCATGGTGTCTGTTCACA | AGTTTTGCTCAGTTTGTCACGTGT |
| chr4 | 134866652 | 134866737 | AAGGAGGAGGGGCCAGGCAGAGTTGAATTGACTTGGGCACACAGACAGACAGATGGACAGACAGACAGACAGACAGACAGGCGGGC | AAGGAGGAGGGGCCAGGCAGAGTTGAATTGACTTGGGCACACAGACAGACAGATGGACAGACAGACAGACAAACGGGT | ACAG,AGG,GACA | B10J | CATGCCTCTCTGACTGTTCTCATC | TTCTGACTGGCCTTGATGGGATTC |
| chr11 | 4263363 | 4263382 | TTTTTGTTTTGTTTTGTTTT | TTTTTGTTTTGTTTTGTTTTGTTTT | TTTTG | B10J | CTTTCTCACTAGGAAGTCAGTGCT | CAAGGAGCTAAAGGGATCTGCA |
| chr9 | 69262368 | 69262407 | CAGGGAGGGAGGCAGGCAGGGAGGCAGGCAGGCAGGCAGG | CAGGGAGGGAGGCAGGCAGGCAGGCAGGCAGG | AGGC | B10J | TGTGAGACCCTGTGTCAATCAC | TTCTGAGCTTTCTAGGTGCTGCAG |
| chr13 | 21034230 | 21034328 | ACAAACCAAACCAAACCAAACCAAACCAAACCAAACCAAACCAAACCAAAACAAAACAACAAAACAAAACAAAACAAAACAAAACAAAACAAAACAAAA | ACAAACCAAACCAAACCAAACCAAACCAAAACAAAACAAAACAAAACAAAACAACAAAACAAAACAAAACAAAACAAAACAAAACAAAA | AACAA,CAAAC | B10J | CAACCTTAGTATTGGAAGATGACGGT | CTGATGACAAAGATTGGACTTGCAG |
| chr18 | 84459108 | 84459170 | GCACACACACACACACACACACAGAGACAGACAGACAGACAGACAGACAGACAGACTGACTGACT | GCACACACACACACACACACACAGAGACAGACAGACAGACAGACAGACAGACTGACTGACT | AGAC,CA,GACT | B10J | AGAATATGCCACCAGACCCAG | GGTTCTCCTAAGCAGAGAGAATGT |
| chr11 | 40320714 | 40320813 | AGCTGCTGCTGTAGAACAAAACCAGCGCTGAGTGCAACCCTTAGAACTGAGAAACTGTGATCCTGCTAGTACCCATCCTTGTGAGAGGGGGGGGGGGGGG | AGCTGCTGCTGTAGAACAAAACCAGCGCTGAGTGCAACCCTTAGAACTGAGAAACTGTGATCCTGCTAGTACCCATCCTTGTGAGAGGGGGGG | GCT,GG | B10J | CAGGTGAGGTGTGGCAACTATG | CTTCTAGGAATGGCTGCATCGG |
| chr14 | 65116731 | 65116777 | CGTGTGTGTGTGTGTGTGTGTGTGTGTGTGTGTGTGTGTGTGTGTGTGT | CGTGTGTGTGTGTGTGTGTGTGTGTGTGTGTGTGTGT | GT | CBA | GATCACATTTATTAAGCAAAAGCTGTGGT | GGACACAGGTTATGCACAGCTTTA |
| chr18 | 31870809 | 31870839 | TCAAACAAACAAACAAACAAAAACAGAAAAG | TCAAACAAACAAATAAACAAACAAAAACAGAAAAA | CAAA | CBA | TCAAAGAGAGTGATGAGAAGGCGAC | CAATGTTGTTGCTGGGACTTGAAC |
| chr5 | 36521268 | 36521292 | CTTGTTTGTTTGTTTGTTTGTTTGT | CTTGTTTGTTTGTTTGTTTGT | TTGT | CBA | TGGGTTGTGGTAATGGTTCTATTCC | TCTGTAGTACCAGAGATATGCACAAATG |
| chr14 | 55280455 | 55280501 | TAATTTACATTATTTTTATGTGTGTGTTTTGTTTGCATGTATGTATA | TTATTACATTATTTTTATGTGTGTGTTTTGTTTGTATG | TG | CBA | TCTTCCACCTTTGATGCTGGGA | CTCGCAGCAGGAAGAGTAAGGAA |
| chr9 | 110027433 | 110027469 | ATTTTTTTTTAATATATATTTATGTTTGCGTGTGCGA | ATATTTTTTTAATATATATTTATGTTTGCGT | TT | CBA | TGCTTGTAACCATGGGTAGGTACAT | AGATGGATGCTGGGAGAAGTGAAG |
| chr8 | 128807001 | 128807063 | AAACCAACCAACCAATCACACACACACACACACACACACACACACACACACACACACACAC | AAACCAACCAACCAATCACACACACACACACACACACACACACACACACACACAC | AACC,CA | CBA | CTTCATAAGTATCCAAGCGTGGTATATAGC | CTGTTGTTTCTGCTGAGTCCTCAC |
| chr11 | 98958183 | 98958224 | TTGTGTGTGTGTGTGTGTGTGTGTGTGTGTGTGTGTGTGCGT | TTGTGTGTGTGTGTGTGTGTGTGCGC | TG | NOD | GAATTTGATGCCCCTTTCCCCAAAG | GGATGTGTGTAGCTCAGTTGGTAGA |
| chr1 | 113169793 | 113171069 | GACCAGAATTCGATAACAAAGAGCTAGAACACTTATACAACATATATTTTTTCTCATATTTTTGGAGGTTGAAAAGTTCCAGCTAAAGCAATAGCAAATCCGGTGAAGACCCTCTTTCTTGTTTGCAGAAAGAGAAAGTTTGGTACATCTTAACATAATAGAGAAGGATAATTTCTCTAGCCTTGTTCATTATATGAACACAAAGTTTGGAGCTGAGACAGAAGGAAGGACCATCCAGAGACTGCCCCAACCAGGGACCCATACCATAATCAGCCACCAAACACAGACACTATTGTATATGCCAGCAAAATTTTGCTGACATGACACTGATATAGCTGTCTCGTGTGAGGCTATGCCAGTGCCTGTCAAATACAAAAATGTATGCTCACAGTCATCTATTGGACAAAATACAGGGCCCCCAATTGAGGAGCTAGAGAAATTACCCAAGGAGCAGAATGGGTCTGTAAACCTATAGGAGGAACAGCGATATGAACTAACCTGTCACCCCTGAGCTCATGTCTCTAGCTGAATATGTAGCAGAAGATGGCCTAATCATCCATCATTGGGAGGAGAGGCCCTTGGTCTTGCAAAGATTATATGCCCCACTTCAGGGGAATGCCAGGGCCAGGAATTAGGAGTGGGTGGCTTGGGGAACAAGGCAAGGGGAGGCTATAGGGAACTTTCCGGATAGTGTTTGAAATGTAAATGAATACGATATCTAATAAAAATTGTTTTAAAAAAAGACTTGGCTGAGATTATGGATTATATAGAATAGATCCTAAATCCATTGAAAAGAGGACTTCTTAATATTCTAAGATTCAGGAATATTTTGAGACTTGTGAGTAATTTAGTTCCTGAATTTAAACATTATTTACTTACTAAATTTCTACAAGTCTCAATTCTGGATGAAATAGTTCAATTAAGAGGGAATTTATAAACAACAGACAGTAGTTGGCATAGCACAGCACTGACAAATTCTGTAACCATGAAGTCTCAAAAATATGGTTCCTAAACAAGGCCTGCAAAATGCAATGCAAGATAGTTTAACAACACAAATGGAAGAACTCTCACTAGGCCCCACAAATTTTTGAAGAACTATAAGCAATTAATGCGTATAGAAAAAGGCAATCAATCTTCTCTAAAGTCAATCACCCTGACAGATTATTCAATCCCAAAAAGTTGCCCCTGCACACATATATAATGATAAATGAATTCTGC | GCCCAGAATTTGATAACAAAGAGCTAGAGCACGTATACAACAAATATTTTTTCTCATATTTCTGGAGGTTGAAAAGTTCCAGCTAAAGCAATAGCAAATCCAGTGAAGACCTTCTTTCTAGTTTGCAGAAAGAGGAAGTTTGGTAAATCTTAACATAATAGAGAAGGATAATTTATCTCTGTGTTCATATATGGTTGCTAAACAAGGCCTGCACAATGCAATGCCAGATAGTATAACAACACAAATGGAAGAACTCTCACTAGGCCAGATAGTATAACAACACAAATGGAAGAACTCTCACTAGACCCCACAAATTTTTGAAGAACTATAAGCAATTAATGGCTATAGAAAAAGGCAGTCAATCTTCTCTCAAGTCCATCACCCTGACAGATTATTCAATCCCAAAAAGTTGCCCCTGCACCACACATATATAATGATAAATGTATTCTGCATTTTGTATCTGTGTGTGTGTGCATGTGTATGTGTGTGTGTGTGTGTGTGTGTGTGTGTGT | TG | NOD | AGTAGGTACAGATTTTAGGACAGTGATC | ACCATTTTCTTTTTAACCTTCAGCTCTG |
| chr4 | 110733622 | 110733910 | CATATATATATAAATATATTACATTATATTACACATATTATTTATATTATATAAATATATTTATATAAGGTGATATATAGTGTAATATATATTATTTATGTTATATAAATAACATATATGTATATATTATATATATATATTGCATCCAGCTCAACCGGCAAGAAAGACGCAACAAACCAGAATCTTCCGCGGCAAGGGCTTTATTGCTTGCTTCTCAGGAAGATCCCAAACCGGGAAAATGGCGCTGCTTTTATAACCTGCAGCGTGACGTTTCAGCACCTGATATGGC | CATATATATATAAATATATTACATTATATTACACATATTATTTATATTATATAAATATATTTATATAAGGTGATATATAGTGTAATATATATTATTTATGTTATATAAATAACATATATGTATATATTATATATATATATATATATTGCATCCAGCTCAACCGGCAAGAAAGACGCAACAAACCAGAATCTTCCGCGGCAAGGGCTTTATTGCTTGCTTCTCAGGAAGATCCCAAACCGGGAAAATGGCGCTGCTTTTATAACCTGCAGCGTGACGTTTCAGCACCTGATATGGT | AT,TA | NOD | CTTCCTAAGTGCACAACTCATTTTAAACC | GCCAGTAATCTCTGAGGAGAGCC |
| chr17 | 4923073 | 4923114 | GAGAGACAGAGACAGAGACAGAGACAGAGACAGAGACAGAGA | GAGAGACAGAGACAGAGAGAGAGAGACAGA | AGAGAC | NOD | GCTAGTCTCAGGAGCAAGGGAG | CAGGTGGGGCAGTGTCATAATC |
| chr5 | 92986933 | 92986967 | ATTTTTTTTTTTTTTAAACAAAGTCTTACTATGTAG | ATTTTTTTTTTTTTTAAACAAAGTCTTA | TT | NOD | TCTGTGGTCTGAGTTATGCATGTGA | CTTTGGTGGAAGGCAGTGCTTAC |
| chr12 | 119494066 | 119494088 | CTTTATTTATTTATTTATTTATT | CTTTATTTATTTATTTATTTATTTATT | TTTA | NOD | CAGGGTGGAAGAGGCAATGATG | GGATGTCATGGAGGGAAATCAAGT |
| chr13 | 109276630 | 109276647 | AATATATATATATATGTG | AATATATATATA | AT | TH | CTTTCCTCTTCAAGGTCAGCAGC | CACCACTGCTGCCAAGATGAG |
| chr1 | 81132699 | 81132743 | CTCTCTGTCTCTGTCTCTGTCTCTGTCTCTCTCTCTCTCTCTC | CTCTCTGTCTCTGTCTCTGTCTCTCTCTCTC | TC,TCTCTG | TH | TGGAAACACACAGGAATCTACAAGG | GTCACTAGATGTTTGTGTAGGGTGTG |
| chr1 | 77561184 | 77561243 | TTATGTATGTATGTATGTGTGTATGTATGTATGTATGTATGTATGTATGTATGTATGTAT | TTATGTATGTATGTATGTGTGTATGTATGTATGTATGTATGTATGTAT | TATG,TGTA | TH | GCCAAACCCATGTGATTCCTAGT | TTAAAGCAGGACTCATGTACTTGTTGC |

| | | | | | | | | |
|---|---|---|---|---|---|---|---|---|
| chr17 | 64827970 | 64828160 | ATGTGTGTGTGTGTATGTATCATATGCCCTTTATCCATTGGGCAGATATCTAAGGTGATCCCACAACTATTGTGAGTTGTGAAGCTATGAGCATGGATGTGCAAGGTTCTCTGTGGTATGTTGACTTAGAATCCTCAGGCATGCGCGCGCGCGCGCGCGCACACACACACACACACACACACACACACA | ATGTGTGTGTGTGTATGTATCATATGCCCTTTATCCATTGGGCAGATATCTAAGGTGATCCCACAACTATTGTGAGTTGTGAAGCTATGAGCATGGATGTGCAAGGTTCTCTGTGGTATGTTGACTTAGAATCCTCAGGCATGCGCACACACACACACACACACACACACA | CA,GC,TG | TH | CAGTTTACATTCAACTAGCACTAGCCTG | ACTCTTCTCTGTAACCTTACCTTGGA |
| chr18 | 55715822 | 55715869 | ATGTGTGTGTGTGTGTGTGTGTGTGTGTGTGTGTGTGTGTGTGTGTGA | ATGTGTGTGTGTGTGTGTGTGTGTGTGTGTGTGTGT | TG | TH | TCTCCTTATATTTCACCTGTGAGACCC | CTTGGAATTCACACAGAGGTCACATG |
| chr15 | 10722050 | 10722069 | TTCTTTTTTTTTTTTTTTTT | TTTTTTTTTTTTTTT | TT | TH | AGTCTGGGCGAATGTTTGTTTCTG | ATGGCATGAACTGTTGACTGTCC |

**Table S5. List of the TRs present in murine homologues of human TR expansion disease genes, related to Figure 1.**

| Gene and Disorder | | | Tandem Repeat (TR) | | | | | | |
|---|---|---|---|---|---|---|---|---|---|
| Human | Disorder | Mouse | Chr | Start | End | Ref | Alt | Location | Strain |
| AFF3 | Intellectual Disability (ID) | Aff3 | 1 | 38248950 | 38248976 | C(TGCTGT)$_3$(TGT)$_2$TG | C(TGCTGT)$_4$TG | exon | FVB,MA/My |
| | | | 1 | 38257463 | 38257487 | C(GCT)$_5$(GCC)$_3$ | C(GCT)$_4$(GCC)$_3$ | exon | SWR |
| AR | Spinal and Bulbar Muscular Atrophy (SBMA) | Ar | X | 97363963 | 97364007 | TA(AA)$_8$G(AA)$_2$G(AA)$_4$GAG(AA)$_5$ | TA(AA)$_8$G(AA)$_2$G(AA)$_4$GAG(AA)$_6$ | 3'UTR | AKR,KK,SMJ |
| ATXN1 | Spinocerebellar Ataxia 1 (SCA1) | Atxn1 | 13 | 45706040 | 45706065 | A(AG)$_6$(AGAA)$_4$A | A(AG)$_8$(AGAA)$_2$A, A(AG)$_7$(AGAA)$_2$A | 3'UTR | AKR,DBA1J,DBA2J,PL,RF |
| | | | 13 | 45707877 | 45707908 | T(TG)$_{15}$T | T(TG)$_{22}$T, T(TG)$_{16}$T, T(TG)$_{20}$T | 3'UTR | AKR,DBA2J,PJ,PL,RF,SEA |
| | | | 13 | 45709443 | 45709466 | A(AAAG)$_3$(AA)$_5$A | A(AAAG)$_3$(AA)$_7$ | 3'UTR | AKR,DBA1J,DBA2J,PL,RF |
| ATXN3 | SCA3<br>Machado-Joseph disease (MJD) | Atxn3 | 12 | 101885459 | 101885478 | C(TT)$_5$(AA)$_4$A | C(TT)$_5$T(AA)$_4$, C(TT)$_4$T(AA)$_5$ | 3'UTR | 129S1,AKR,BALB,BTBR,BUB,C57L,C58J,CBA,CE,FVB,ILNJ2,KK,LP,MA/My,NOD,NOR,NZB,NZO,NZW,PJ,RF,SEA,SJL,SMJ,SWR,TH |
| | | | 12 | 101885752 | 101885792 | G(CA)$_6$AAAGTATTCCTCAAATT(AA)$_5$A | G(CA)$_6$AAAGTATTCCTCAAATT(AA)$_5$ | 3'UTR | 129S1,AKR,BALB,BTBR,BUB,C57L,C58J,CBA,DBA1J,DBA2J,FVB,ILNJ2,KK,LP,MA/My,NOD,NOR,NZB,NZO,NZW,PJ,PL,RF,SEA,SJL,SMJ,SWR,TH |
| ATXN7 | SCA7 | Atxn7 | 14 | 8363465 | 8363524 | T(TC)$_5$(CT)$_4$CAT(AC)$_{19}$ | T(TC)$_8$(CT)$_4$CATACAT(AC)$_{21}$, T(TC)$_8$(CT)$_4$CATACAT(AC)$_{22}$, T(TC)$_8$(CT)$_4$CAT(AC)$_{27}$, T(TC)$_8$(CT)$_4$CAT(AC)$_{25}$ | 3'UTR | AJ,AKR,BALB,BTBR,BUB,C3H,C58J,DBA2J,ILNJ2,NZB,NZO,NZW,RF,SMJ |
| | | | 14 | 8363981 | 8364009 | C(TT)$_2$A(TTTTATT)$_3$TT | C(TTTTATT)$_3$TT | 3'UTR | 129S1,C3H,C57L,CBA,DBA1J,DBA2J,KK,MRL,NZW,PJ,PL |
| BEAN1 | SCA31 | Bean1 | 8 | 104944867 | 104944925 | A(AC)$_5$ATGTA(CC)$_2$AC(AA)$_{12}$GC(AA)$_5$C | A(AC)$_5$ATGTA(CC)$_2$AC(AA)$_5$A, A(AC)$_5$ATGTA(CC)$_2$AC(AA)$_6$ | 3'UTR | KK,PL |
| CACNA1A | SCA6 | Cacna1a | 8 | 85365341 | 85365372 | A(CAC)$_6$(CAT)$_4$C | A(CAC)$_7$(CAT)$_4$C | exon | AJ,C3H,CBA,CE,DBA1J,DBA2J,FVB,ILNJ2,KK,MA/My,NOD,NOR,NZW,RH2,SJL,SMJ,SWR |
| COMP | Multiple Epiphyseal Dysplasia 1 (EDM1)<br>Pseudoachondroplasia (PSACH) | Comp | 8 | 70834622 | 70834645 | ACAAGCTACGG(GT)$_5$GGG | ATAAGCTACGG(GT)$_7$ | 3'UTR | 129S1,AKR,BALB,BUB,C3H,C57L,C58J,CBA,CE,DBA1J,DBA2J,FVB,ILNJ2,KK,MA/My,MRL,NZB,NZO,NZW,PL,RF,SJL,SMJ,SWR,TH |
| DAB1 | SCA37 | Dab1 | 4 | 104601400 | 104601420 | C(TGGTTT)$_3$TG | C(TGGTTT)$_2$TG(TT)$_2$TG, CTGGTTT(TGTTTT)$_2$TG | 3'UTR | AJ,AKR,BALB,C3H,C58J,CBA,CE,ILNJ2,KK,MA/My,MRL,NOD,NOR,PJ,PL,RF,SEA,SJL,SWR,TH |
| DIP2B | ID | Dip2b | 15 | 99936546 | 99936567 | T(GGC)$_7$ | T(GGC)$_5$ | 5'UTR | 129S1,C57L,ILNJ2,NZW,PJ,SEA |
| | | | 15 | 100115041 | 100115079 | C(TT)$_5$CT(TTTC)$_3$(TT)$_7$ | C(TT)$_5$CT(TTTC)$_3$(TT)$_7$T, C(TT)$_4$TCT(TTTC)$_2$(TT)$_3$C(TT)$_4$ | 3'UTR | AJ,SMJ |
| | | | 15 | 100116410 | 100116462 | TC(GT)$_{25}$G | TC(GT)$_{23}$G, TC(GT)$_{24}$G, TC(GT)$_{21}$G, TC(GT)$_{19}$G | 3'UTR | 129S1,AJ,B10J,BALB,BTBR,BUB,C3H,C57L,C58J,CBA,CE,DBA1J,DBA2J,ILNJ2,MA/My,MRL,NOR,NZO,NZW,PL,SJL,SMJ,SWR |
| DMD | Duchenne Muscular Dystrophy (DMD) | Dmd | X | 84246546 | 84246583 | G(CA)$_{17}$CGC | G(CA)$_{13}$CGC | 3'UTR | CE |
| DMPK | Myotonic Dystrophy 1 (DM1) | Dmpk | 7 | 18827113 | 18827133 | TCGCG(CC)$_8$ | TCG(CC)$_7$, TCG(CC)$_7$C, TCGCG(CC)$_7$C, TCGCG(CC)$_7$, TCG(CC)$_9$, TCG(CC)$_9$C | 3'UTR | 129S1,AKR,B10J,BTBR,BUB,C57L,CE,MA/My,NOD,NZB,NZO,PL,SJL,SWR |
| FMR1 | Fragile X-associated Disease | Fmr1 | X | 67722328 | 67722349 | CG(CGG)$_6$CG | CG(CGG)$_8$CG | 5'UTR | ILNJ2 |
| FOXL2 | Blepharophimosis, Ptosis and Epicanthus Inversus (BPES) | Foxl2 | 9 | 98838553 | 98838575 | A(CCG)$_3$CCA(CCG)$_3$C | A(CCG)$_5$C, A(CCG)$_7$C | exon | CE,ILNJ2,NZO,NZW,PJ,SEA |
| FXN | Friedreich Ataxia (FRDA) | Fxn | 19 | 24238851 | 24238883 | A(CTTT)$_4$(TT)$_7$TC | A(CTTT)$_4$(TT)$_9$T, A(CTTT)$_4$(TT)$_9$ | 3'UTR | C3H,PJ,SWR |
| GLS | Global Developmental Delay, Progressive Ataxia and Elevated Glutamine (GDPAG) | Gls | 1 | 52203756 | 52203800 | C(GT)$_{22}$ | C(GT)$_{21}$, C(GT)$_{24}$, C(GT)$_{25}$, CAT(GT)$_2$ATGA(GT)$_{30}$ | 3'UTR | BALB,BTBR,BUB,C57L,C58J,MA/My,MRL,NZO,NZW,PL,RF,RH2,SEA,SMJ,SWR |
| | | | 1 | 52204131 | 52204177 | AAACAA(CAACC)$_3$(AAACC)$_2$(AAACA)$_3$A | AAACAA(CAACC)$_3$(AAACC)$_3$(AAACA)$_2$A | 3'UTR | BTBR,C57L,KK |
| | | | 1 | 52223535 | 52223588 | T(TC)$_{18}$T(CA)$_8$ | T(TC)$_{20}$T(CA)$_9$, T(TC)$_{28}$T(CA)$_6$TA, T(TC)$_{19}$T(CA)$_8$, T(TC)$_{21}$T(CA)$_{16}$ | 3'UTR | AKR,BALB,C57L,NOD,NZW,PL,SJL |
| | | | 1 | 52271853 | 52271881 | G(GCT)$_9$G | G(GCT)$_6$G, G(GCT)$_8$G, G(GCT)$_7$G | exon | BTBR,C57L,KK,NZO,NZW,SMJ,SWR |
| | | | 1 | 52272301 | 52272329 | A(CGC)$_3$CCT(CGC)$_5$C | A(CGC)$_3$CCT(CGC)$_6$C, A(CGC)$_3$CCT(CGC)$_7$C | 5'UTR | BTBR,C57L,KK,SWR |
| HTT | Huntington Disease (HD) | Htt | 5 | 34919310 | 34919387 | A(CAG)$_4$CCA(CCG)$_2$CAGG(CGC)$_3$CAC(CGC)$_4$CTCCGCCTCAA(CC)$_2$CTCA(GCC)$_3$ | A(CAG)$_4$CCA(CCG)$_2$CAGG(CGC)$_3$CAC(CGC)$_3$CTCCGCCTCAA(CC)$_2$CTCA(GCC)$_3$ | exon | BUB,C58J,FVB,MA/My,MRL,NOD,NOR,NZB,PL,SWR,TH |
| | | | 5 | 35069244 | 35069303 | CT(TCCC)$_2$TCCT(TCCC)$_6$TCCG(TCCC)$_4$TC | CC(TCCC)$_9$TCCG(TCCC)$_4$TC, CC(TCCC)$_8$TCCG(TCCC)$_4$TC | 3'UTR | BUB,C58J,DBA1J,DBA2J,FVB,MA/My,MRL,NOD,NOR,NZB,PL,SWR,TH |
| JPH3 | Huntington Disease-Like 2 (HDL2) | Jph3 | 8 | 122457253 | 122457326 | A(AGCC)$_3$AACCAAGACTCCGGTTTGTTGATCGCCTTGTTTC(TT)$_{13}$T | A(AGCC)$_3$AACCAAGACTCCGGTTTGTTGATCGCCTTGTTTC(TT)$_{13}$ | 5'UTR | SEA,SMJ |
| NIPA1 | Amyotrophic Lateral Sclerosis (ALS) | Nipa1 | 7 | 55627999 | 55628023 | T(TTTTG)$_4$(TT)$_2$ | T(TTTTG)$_6$(TT)$_2$, T(TTTTG)$_5$(TT)$_2$, T(TTTTG)$_7$(TT)$_2$, T(TTTTG)$_8$(TT)$_2$ | 3'UTR | 129S1,AJ,AKR,BALB,BTBR,BUB,C3H,C57L,CBA,CE,DBA1J,DBA2J,FVB,KK,MA/My,MRL,NOD,NOR,NZB,NZO,PL,RF,SEA,SMJ,SWR,TH |
| PABPN1 | Oculopharyngeal Muscular Dystrophy (OPMD) | Pabpn1 | 14 | 55131614 | 55131644 | T(GGC)$_7$(AGC)$_3$ | T(GGC)$_2$AGC(GGC)$_4$(AGC)$_3$ | exon | NZB |
| | | | 14 | 55135561 | 55135588 | G(AA)$_5$GAATT(AA)$_6$ | G(AA)$_5$AGAATT(AA)$_6$, G(AA)$_5$GAATT(AA)$_6$A | 3'UTR | DBA2J,SMJ |
| PPP2R2B | SCA12 | Ppp2r2b | 18 | 42778353 | 42778372 | A(TT)$_5$(TTG)$_3$ | A(TT)$_4$T(TTG)$_3$, A(TT)$_2$T(TTG)$_4$ | 3'UTR | 129S1,AJ,AKR,BALB,C3H,C57L,C58J,CBA,DBA1J,DBA2J,FVB,ILNJ2,KK,MRL,NZB,NZO,NZW,PJ,PL,RF,RH2,SEA,SJL,SMJ,SWR,TH |
| PRDM12 | Hereditary Sensory and Autonomic Neuropathy 8 (HSAN8) | Prdm12 | 2 | 31545498 | 31545538 | AAT(AC)$_{19}$ | AAT(AC)$_{20}$, AAT(AC)$_{17}$ | 3'UTR | MA/My,NZO |
| RAPGEF2 | Familial Adult Myoclonic Epilepsy 7 (FAME7) | Rapgef2 | 3 | 78970854 | 78970907 | T(AA)$_2$(AAAAAAAC)$_3$CC(AA)$_2$AC(AA)$_2$(AAAC)$_3$A | T(AA)$_3$(AAAAAAAC)$_2$(AA)$_3$CCC(AA)$_2$AC(AA)$_2$(AAAC)$_3$A, T(AA)$_3$A(AAAAAAAC)$_2$(AA)$_3$CCC(AA)$_2$AC(AA)$_2$(AAAC)$_3$A | 3'UTR | BUB,C57L,CE,ILNJ2,MRL,NZB,RH2,SEA,SWR,TH |
| RUNX2 | Cleidocranial Dysplasia 1 (CLCD1) | Runx2 | 17 | 44915737 | 44915759 | GG(TT)$_4$TG(TT)$_5$T | GG(TT)$_4$TG(TT)$_6$T, G(TT)$_6$G(TT)$_5$, G(TT)$_5$(GTT)$_4$(TT)$_2$, GG(TT)$_4$TG(TT)$_5$ | 3'UTR | BTBR,C57L,FVB,NZW,SJL,SMJ,SWR |
